# HiTIPS: High-Throughput Image Processing Software for the Study of Nuclear Architecture and Gene Expression

**DOI:** 10.1101/2023.11.02.565366

**Authors:** Adib Keikhosravi, Faisal Almansour, Christopher H. Bohrer, Nadezda A. Fursova, Krishnendu Guin, Varun Sood, Tom Misteli, Daniel R. Larson, Gianluca Pegoraro

## Abstract

High-throughput imaging (HTI) generates complex imaging datasets from a large number of experimental perturbations. Commercial HTI software for image analysis workflows does not allow full customization and adoption of new image processing algorithms in the analysis modules. While open-source HTI analysis platforms provide individual modules in the workflow, like nuclei segmentation, spot detection, or cell tracking, they are often limited in integrating novel analysis modules or algorithms. Here, we introduce the High-Throughput Image Processing Software (HiTIPS) to expand the range and customization of existing HTI analysis capabilities. HiTIPS incorporates advanced image processing and machine learning algorithms for automated cell and nuclei segmentation, spot signal detection, nucleus tracking, spot tracking, and quantification of spot signal intensity. Furthermore, HiTIPS features a graphical user interface that is open to integration of new algorithms for existing analysis pipelines and to adding new analysis pipelines through separate plugins. To demonstrate the utility of HiTIPS, we present three examples of image analysis workflows for high-throughput DNA FISH, immunofluorescence (IF), and live-cell imaging of transcription in single cells. Altogether, we demonstrate that HiTIPS is a user-friendly, flexible, and open-source HTI analysis platform for a variety of cell biology applications.

## Introduction

High-Throughput Imaging (HTI) fully automates the acquisition and analysis of large fluorescence microscopy imaging datasets. HTI was originally developed to provide phenotypic readouts for large high-throughput chemical screens to identify compounds with desirable therapeutic activities ^1,2^. Since then, this technology has been widely adapted to work in conjunction with functional genomics screens to identify molecular pathways involved in a variety of cellular functions. HTI has also been extensively used in more traditional cell biology applications, where the automation of image acquisition and analysis has been used to systematically quantify at the single-cell level events that are heterogeneous, rare, or dynamic in cellular populations ^2^.

The study of nuclear architecture and gene expression has particularly benefited from HTI. DNA Fluorescence In Situ Hybridization (FISH)-based HTI imaging has been used to study the spatial organization of genes and chromosomes within the nucleus, providing insights into the mechanisms of gene regulation and nuclear architecture. For example, FISH-based high-throughput imaging has been used to study the three-dimensional organization of the genome ^3–7^. In addition, IF-based HTI assays have been used to probe nuclear architecture using fluorescently labelled endogenous architectural markers in combination with functional genomics screens ^8–13^. Finally, HTI has also been used to explore the dynamics of transcription initiation and RNA splicing at the single-cell level ^14,15^.

HTI has been enabled by the development of automated microscopy platforms and image analysis software to rapidly acquire and process large amounts of fluorescence microscopy data. These tools allow the extraction of quantitative information at the single-cell level of up to hundreds of cellular features in individual cells ^16^. Traditionally, software for HTI analysis has three major components: 1) individual analysis modules to perform basic HTI analysis steps (e.g., nuclear segmentation, spot detection, cell tracking, fluorescence intensity measurements), 2) a graphical user interface (GUI) for end-users without programming skills to set the parameters for the analysis modules in an interactive fashion, and 3) a mechanism to chain the analysis modules into end-to-end analysis pipelines that can run in batch mode. Previous software tools, such as FISH-quant ^17^ and CellProfiler ^18^, have been introduced to tackle the wide demand for cellular and FISH image analysis. However, while covering an extremely large share of both traditional and advanced automated image analysis cases, some complex cellular features have been refractory to analysis, such as measurement of physical distance between genomic loci or transcription sites ^4^, measurement of clustering of centromeres ^3^, and detection, registration, and tracking of transcription sites in the nucleus of live cells ^14,15^. Furthermore, for many analysis packages users often face challenges including a steep learning curve, the absence of real-time feedback, and the necessity of complex configuration and optimization to initiate an analysis. For some analysis packages, users also need a degree of programming knowledge for the creation of analysis pipelines and customization of the analysis workflow.

We developed a software platform for HTI analysis, which we named High-Throughput Image Processing Software (HiTIPS), as a multi-application image analysis pipeline for a wide range of HTI imaging assays processes. HiTIPS is open source, built in Python, and provides a graphical user interface (GUI) for interactive and user-friendly visualization of images, and for the optimization of HTI analysis parameter settings without the need for coding. Furthermore, the GUI provides access to a variety of image analysis modules for HTI, which incorporate both traditional image processing algorithms and advanced machine learning algorithms for nucleus segmentation and fluorescence signal spot detection. In addition, HiTIPS includes advanced methods for tracking and registration of nuclei of live cells in timelapse experiments. Importantly, HiTIPS is built using a flexible architecture, which allows the incorporation of novel algorithms, and the addition of new plugins for additional HTI analysis tasks, making HiTIPS extensible and amenable to a wide variety of HTI assays.

## Results

### Image Loading and Visualization

HiTIPS is designed for easy and versatile setup of HTI image analysis pipelines. This is achieved by first optimizing analysis parameters in an interactive fashion on a subset of representative images. This process is achieved by providing the user visual feedback of the results of the analysis overlayed on the original images. Once this optimization process is completed, the user can then run the analysis on the whole dataset in batch mode. Both the interactive analysis module setup and launching the analysis pipelines steps in HiTIPS do not require programming, thanks to a graphical user interface (GUI) that is used for data loading, for image and results visualization, and for the choice of the image analysis modules parameters (Fig. 1A).

**Fig. 1).**
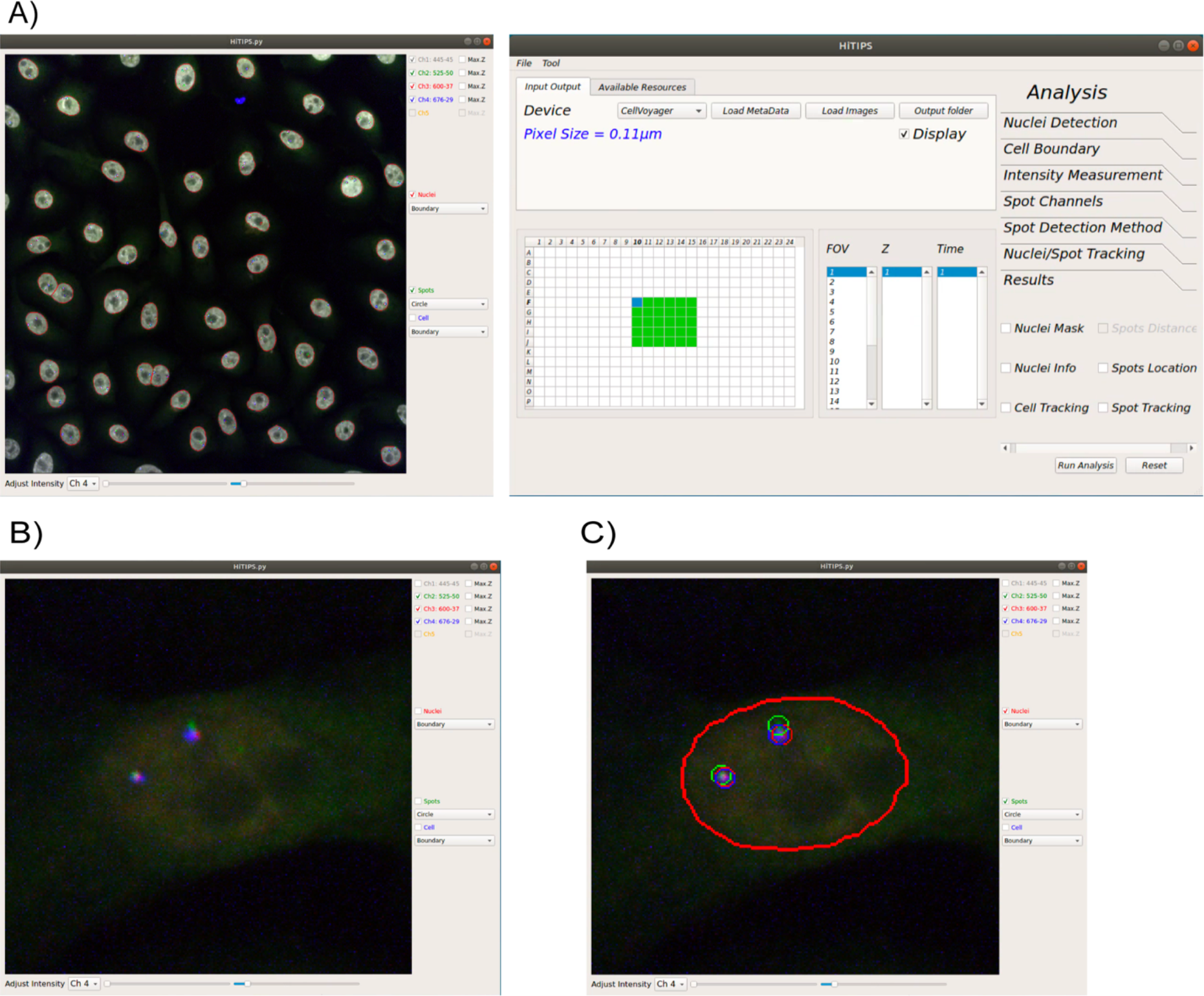
Examples of the HiTIPS Graphical User Interface (GUI). **A)** Representative screenshot of the HiTIPS GUI for HTI dataset selection and on-demand image loading, with metadata loading and integration in various formats, including CellVoyager and Micro-Manager. **B)** Representative screenshot of the image visualization controls in the HiTIPS GUI, including, among others, fluorescence channel toggling and z-projected views for 3D z-stacks, and fluorescence intensity visualization adjustment. **C)** Representative screenshot of the overlayed display in the HiTIPS GUI of nuclei masks borders (red) and spot detection in 2 different channels (red and green circles) output from image HiTIPS analysis modules.

HiTIPS allows users to select HTI imaging datasets and to load data on demand, thus eliminating the need to retain the entire dataset in memory (Fig. 1A). This enables swift, effective access to extensive image datasets, while minimizing memory requirements for processing. In addition, HiTIPS uses either a generic Bio-Formats reader ^19^, which allows the loading and conversion of more than 120 different imaging formats, or it uses image acquisition metadata (Well position, field of view (FOV), channel, etc.) automatically generated by the microscope and saved in separated XML files. While this second mechanism is currently only implemented for the CellVoyager format from Yokogawa and for imaging datasets generated by Micro-Manager, the open source and modular nature of HiTIPS allows the future extension of metadata reading from files to other instruments and formats, potentially including the recently developed OMERO-ZARR format ^20^. Thanks to the use of image acquisition metadata by HiTIPS, users can select specific wells, FOVs, and/or channels to quickly load single merged FOVs in the viewer for visual inspection and for optimization of the image analysis parameters (Fig. 1A).

Once the images are loaded, users can perform a series of routine changes to their visualization, including toggling specific channels on or off, showing a z-projected version of the image if the FOV is present as a 3D z-stack, and independently adjusting minimum and maximum intensity levels for each of the channels (Fig. 1B). Visual inspection of random wells and FOVs in the dataset is often an essential quality control step before setting up an HTI image analysis pipeline, and it is greatly facilitated by rapid loading and rendering of the images by HiTIPS. Furthermore, the image visualization interface is not limited to the original images, but also includes the overlayed presentation of object masks and borders generated by different image analysis modules that can also be selected and whose parameters can be modified using another window in the GUI (Fig. 1C). This is an essential feature that enables rapid cycles of parameter optimization during the interactive image analysis setup phase. Finally, after configuring the analysis parameters in the interactive mode, HiTIPS allows users to choose the number of parallel processing threads for batch analysis depending on the technical specification of the hardware on which the application is running.

### HTI Image Analysis Workflow

While HiTIPS was built to analyze a variety of HTI assays, and it can further be extended or customized by a developer to accommodate additional specific analysis needs, we used the study of genome architecture and gene expression in both fixed and live cells as a model system for our initial efforts in the development of this software. For this reason, the HiTIPS analysis workflow currently includes sequential use of image and metadata loading (Fig. 2, i), nuclei segmentation (Fig. 2, ii), fluorescent spot finding (Fig. 2, iii), nuclei tracking (Fig. 2, iv), nuclei and spot patch generation (Fig. 2, v), nuclei and spot patch registration (Fig. 2, vi), spot assignment to a track (Fig. 2, vii), measurement of fluorescence intensity (Fig. 2, viii), and 2-state Hidden Markov Model (HMM) fitting to segment fluorescence intensity traces (Fig. 2, ix). HiTIPS also allows the selection of the workflow steps only up to the spot finding module (Fig. 2,i – ii), or up to the nucleus tracking module for live-cell HTI assays that do not require spot level measurements (Fig. 2, i, ii, and iv). This selection can be performed by toggling specific modules on or off in the GUI (Fig. 1A) during the interactive setup phase of image analysis workflow.

**Fig. 2).**
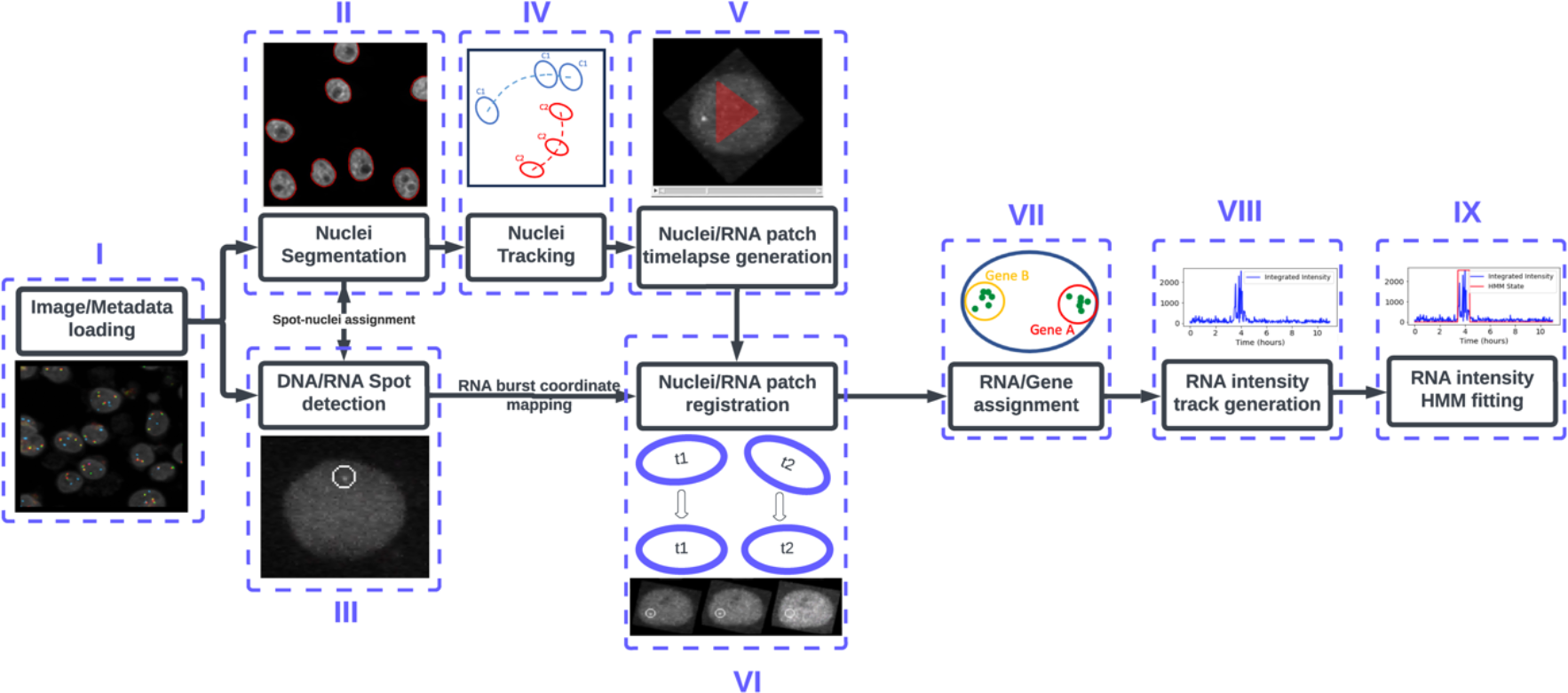
Schematic representation of the full HiTIPS image analysis workflow. i) Image and metadata loading, leading to ii) Nuclear segmentation and iii) Spot finding, iv) Nucleus tracking v) Single nucleus timelapse generation, and vi) Frame to Frame nuclei registration processes. vii) Spot assignments to specific tracks are determined before viii) Measurement of track fluorescence intensities, culminating in ix) Segmentation of gene ON and OFF states by fitting a 2-state Hidden Markov Model (HMM) to the fluorescence intensity tracks.

To show the utility of HiTIPS across a broad spectrum of data types and applications related to the biology of the cell nucleus, we applied it to three distinct HTI assays: 1) measurement of 3D physical distances between two genomic loci visualized with DNA FISH probes, 2) estimation of spatial clustering in the nucleus of centromeres labelled in IF with an antibody to the centromeric protein CENPC, and 3) a set of high-throughput measurements of transcriptional activity in live cells of the endogenous *KPNB1* or *ERRFI1* genes labelled with the fluorescent MS2/MCP-GFP system.

### HiTIPS measures genomic locus to locus distances in high-throughput fashion

Mammalian genomes are spatially organized in the cell nucleus at several different hierarchical levels, and 3D genome organization is tightly correlated with many nuclear functions such as transcription, replication, and DNA damage repair (Misteli 2020). A prominent feature of genome organization are Topologically Associating Domains (TADs), which represent genomic regions which exhibit an enhanced propensity for mutual interaction, across relatively large genomic distances (200 kb - 1 Mb) ^21,22^. At the molecular level, one of the key factors for TAD establishment and maintenance is the cohesin complex, as demonstrated by the observation that acute depletion of the RAD21 cohesin subunit leads to complete disappearance of these domains as measured by biochemical chromatin conformation capture techniques ^23^, and to an increase in physical distances between adjacent TADs as measured by DNA FISH imaging ^24^.

We wanted to benchmark HiTIPS in a high-throughput DNA FISH assay by testing whether we could measure changes in the physical distances of the boundaries of the TAD containing the human *EGFR* gene on Chr 7 upon acute depletion of RAD21 (Fig. 3A). To this end, we performed high-throughput DNA FISH imaging experiments in HCT116-RAD21-AID cells, where RAD21 can be rapidly degraded by the cellular ubiquitin/proteasome machinery upon binding of the AID degron domain to Auxin ^25^ (Fig. 3A). We used automated confocal imaging to acquire z-stack images of HCT116-RAD21-AID treated for 3 hrs with Auxin or mock treated cells in 3 channels (DAPI, EGFR TAD 5’ boundary/Probe A, EGFR 3’ boundary/Probe B, Fig. 3B). 3D image stacks were analyzed with HiTIPS in batch by segmenting nuclei using the DAPI image, finding the position of FISH spot in 3D, and by calculating minimum distances between FISH spot centers in the two different channels on a per allele fashion (Fig. 2, i – iii). By plotting the distribution of minimum distances between the genomic loci at the base of the loop domain in 1874 cells in either Auxin-treated or mock-treated control cells, we observed that RAD21 degradation upon Auxin treatment led to a statistically significant increase in the distance between TAD boundaries (Fig. 3C, p < 2e-16, Wilcoxon Test). These results show that HiTIPS can be used for the automated analysis of 3D distances measured from DNA FISH images.

**Fig. 3).**
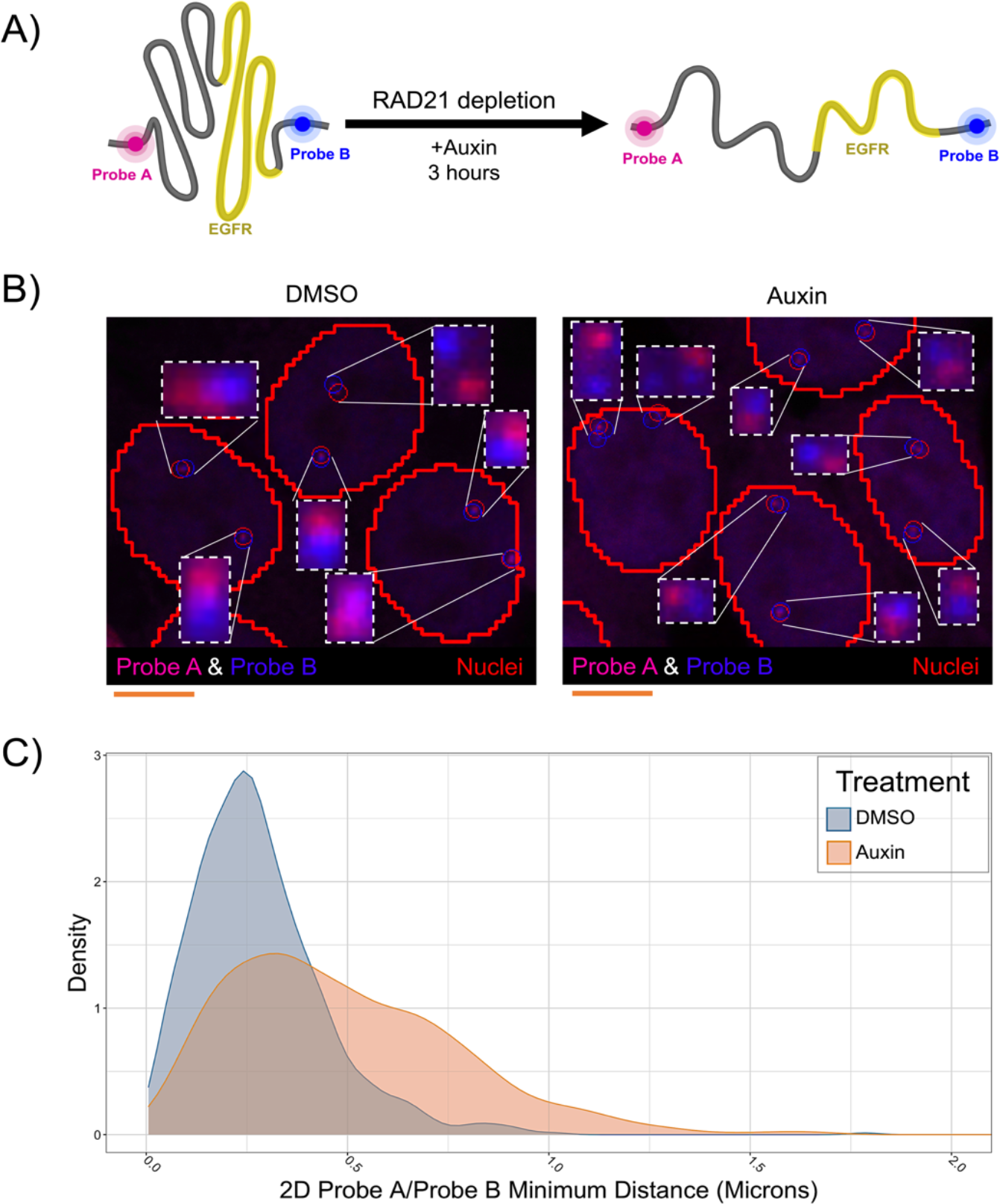
HiTIPS measures distances between genomic loci labelled with High-Throughput DNA FISH probes. **A)** Schematic representation of the experiment to measure spatial distances of two genomic loci located at the base of a TAD encompassing the *EGFR* gene on Chr 7 and detected by FISH probes in different channels (Probe A and Probe B). Treatment with Auxin in HCT116-RAD21-AID cells leads to rapid proteolytic degradation of the RAD21-AID fusion protein via the ubiquitin/proteasome pathway. **B)** Representative 3D maximally projected images of HCT116-RAD21-AID cells stained with DNA FISH probes A and B targeted to the 5’ and 3’ boundaries of the EGFR TAD, respectively, and of the results of the HiTIPS spot finding algorithm results overlayed as red and blue circles, respectively. Scale bar: 5 microns. **C)** Density plots of minimum spatial distances between the A and B DNA FISH probes in 1874 cells treated with Auxin or DMSO.

### Clustering analysis of centromeric chromosomal regions in the nucleus

Centromeres are specialized genomic regions that assemble the kinetochore, a large protein complex consisting of several components including the evolutionarily conserved CENPC protein ^26^. Kinetochores physically connect chromosomes to microtubules and ensure high-fidelity genome segregation during cell division ^27^. Centromere positions within the 3D space of cell nucleus vary across species ^28^. Recently, it was shown that loss of the condensin II complex subunit NCAPH2 leads to centromere clustering in human cancer cells using biochemical techniques and traditional low-throughput fluorescence microscopy ^29^.

We tested whether we could use HiTIPS to measure spatial clustering of centromeres at the single-nucleus level (Fig. 4). To this end, we reverse-transfected HCT116-Cas9 cells with siRNA oligos against the *NCAPH2* gene, or a scrambled negative siRNA control in 384-well plates. Cells were fixed and stained with DAPI for nuclei segmentation and with a CENPC-specific antibody to visualize centromeres. Stained cells were then imaged in 3D (Fig. 4A), and maximally projected images were analyzed using HiTIPS for nuclear segmentation and spot finding/localization (Fig. 2, i – iii). HiTIPS was capable to precisely detect and localize CENPC spots in cell nuclei, even in regions of high density of CENPC spots (Fig. 4A). More importantly, the single CENPC spots position datasets and the nuclei ROIs could be used in a separate analysis to calculate a centromeric clustering score in single cells. We defined this score as the percentage of the measured curve for Ripley’s K function ^30^ that is above the curve for the random Poisson distribution (See Materials and Methods for details). This clustering analysis showed that, in line with visual inspection, cells transfected with siNCAPH2 had higher clustering scores (Fig. 4B) and fewer distinct centromeres (Fig. 4C) than cells transfected with the control scrambled siRNA. These results show the utility of HiTIPS in the analysis of IF-based HTI assays at the single-cell level, and they confirm previous traditional fluorescence microscopy-based results ^29^, while expanding the phenotypic analysis from a few cells to thousands of cells for each condition.

**Fig. 4).**
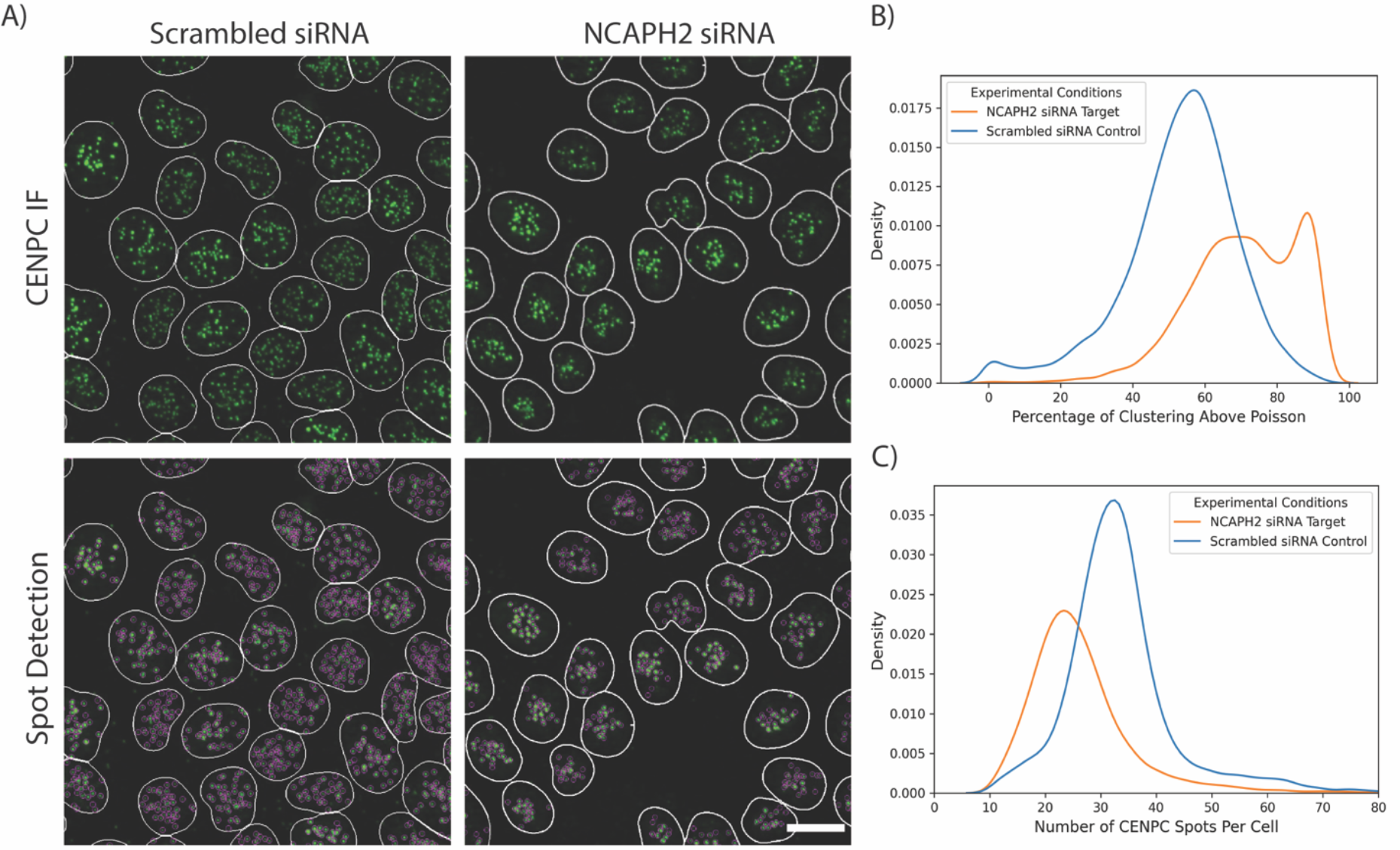
HiTIPS analysis of centromeric clustering using. **A)** 3D maximally projected images of HCT116-Cas9 cells reverse transfected in 384-well plates for 72 hrs with either a scrambled non-targeting siRNA (siScramble), or with an siRNA against the condensin II subunit NCAPH2 (siNCAPH2). Transfected cells were stained in IF with DAPI and a CENPC antibody, and imaged using a high-throughput confocal spinning disk microscope. The border of the segmented nuclei masks is overlayed on the image and colored in white. CENPC spots detected by HiTIPS are overlayed as magenta circles in the lower images. Scale bar: 10 microns. **B)** Density plot of cell-level CENPC spots clustering scores for siScramble and siNCAPH2 as calculated by HiTIPS. Higher values of the clustering score indicate more clustering of CENPC spots in the nucleus. **C)** Density plot of the number of CENPC spots per cell for siScramble and siNCAPH2 as calculated by HiTIPS.

### Semi-automated Measurement of Transcription Dynamics at the Single-Allele Level in Live Cells

Random or targeted intronic integration of endogenous genes with arrays of MS2 hairpins in cells stably expressing a fluorescently tagged fusion of the MS2 capsid protein (MCP) has been instrumental in demonstrating that in mammalian cells transcription happens in bursts of activity followed by periods of inactivity ^31^, and that splicing of long introns is recursive ^15^. HTI acquisition and analysis has been used to measure the dynamics of these events in large numbers of single live cells ^14,15^.

We aimed to compare HiTIPS with our previous image analysis pipeline ^15^ to show that it can be applied to precisely quantify transcriptional dynamics in live cells at the single-cell level (Fig. 5). To this end, we ran the full HiTIPS analysis pipeline on HTI datasets from two clonal human bronchial epithelial (HBEC) cell lines, in which the *ERRFI1* and *KPNB1* genes have been endogenously tagged with 24xMS2 loops, enabling visualization of their nascent transcription with MCP-GFP ^15^. MS2/MCP-GFP labelled transcription sites in these nuclei were detected (Fig. 2, iii), registered (Fig. 2, vi), grouped into tracks (Fig. 2,vii), and the integrated fluorescence intensity was measured at the site of transcription (Fig. 2, viii).

**Fig. 5.**
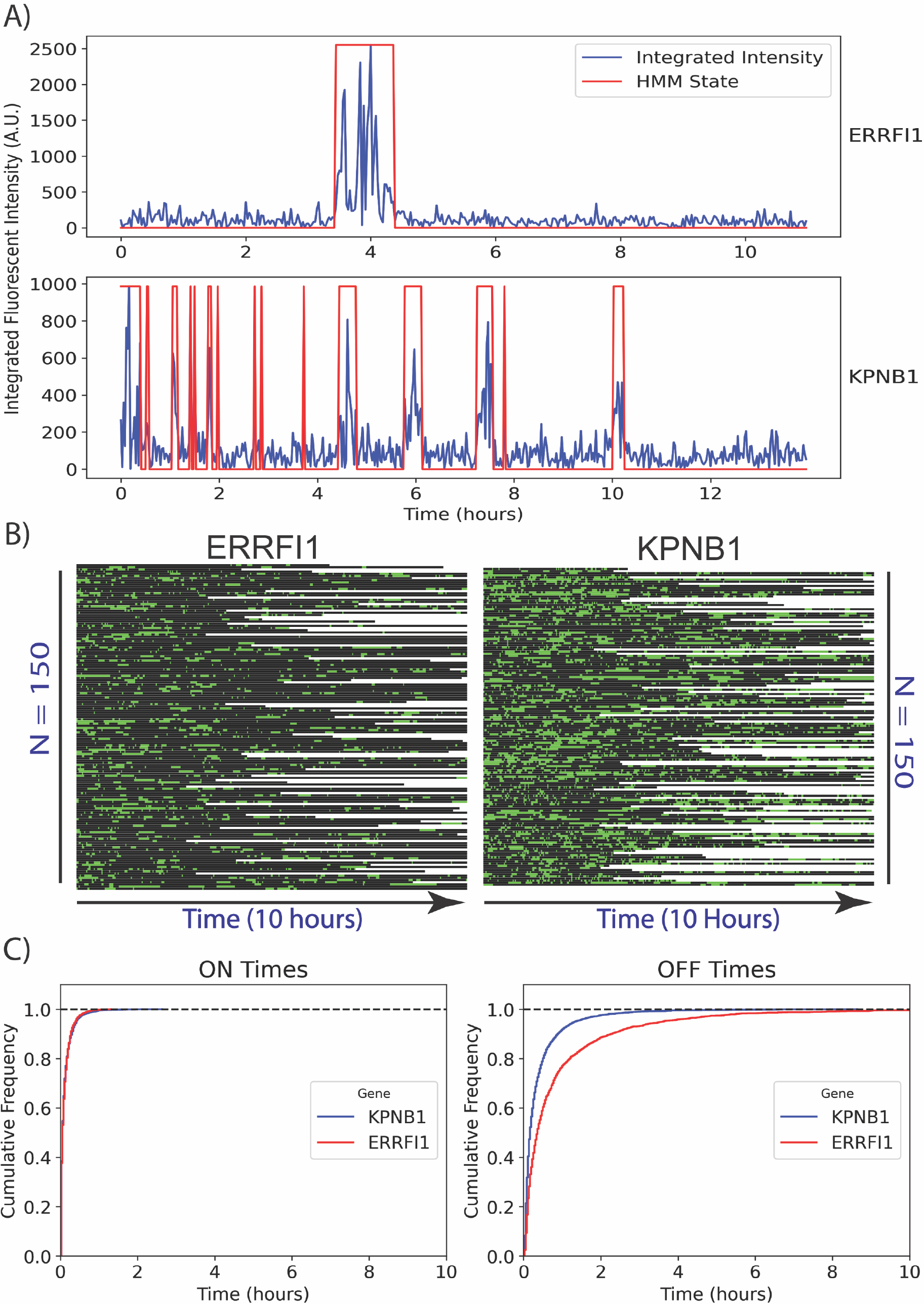
HiTIPS quantifies gene expression dynamics in live cells. **A)** Normalized and integrated fluorescence intensity plot at the site of MS2/MCP-GFP spots detected in live cells over 10 hours for both MS2-tagged *ERRFI1* and *KPNB1* genes. The HMM 2-state segmentation of intensity traces into “ON” and “OFF” times is indicated in red. **B)** Kymograph of 2-state HMM-segmented fluorescence intensity traces for a subsample of fluorescence intensity traces (n = 150 for ERRFI1-MS2/MCP-GFP and n = 150 for KPNB1-MS2/MCPGFP) over 10 hours. Green represents “ON” times (active transcription), black represents “OFF” times (no transcription) in the kymograph. **C)** Empirical cumulative distribution function plots of ON and OFF time durations for both cell lines across multiple cells and transcription sites (n = 260 for KPNB1-24xMS2/MCP-GFP and n = 260 cells for both ERRFI1-24xMS2/MCPGFP).

This automated analysis resulted in the generation of 1232 fluorescence intensity traces (690 for ERRFI1-24xMS2/MCP-GFP and 542 for KPNB1-24xMS2/MCP-GFP). We further conducted a visual quality control step on the segmented transcription site traces to obtain a total of 595 traces (277 for ERRFI1-24xMS2/MCP-GFP and 318 for KPNB1-24xMS2/MCP-GFP). In agreement with previous work ^15^, visualization of a subset of fluorescence intensity traces for single alleles of *ERRFI1* and *KPN1B1* revealed that *KPNB1* bursts more frequently than *ERRFI1*, due to shorter periods of inactivity (OFF times) (Fig. 5A). Extending this analysis to a larger subsample of fluorescent intensity traces for each cell line (Fig. 5B, n = 150 for ERRFI1-24xMS2/MCP-GFP, and n = 150 for KPNB1-24xMS2/MCP-GFP) confirmed that the observed difference in transcriptional dynamics of *KPNB1* and *ERRFI1* extends to the cellular population. Further evidence supporting this observation comes from analysis of cumulative distributions of ON and OFF times across an even larger subsample of all the intensity traces (Fig. 5C, n = 260 for ERRFI1-24xMS2/MCP-GFP, and n = 260 for KPNB1-24xMS2/MCP-GFP). We observed that *ERRFI1* has on average significantly longer OFF times compared to *KPNB1*, while the distribution of ON times is not substantially different between the two genes. These results obtained with HiTIPS pipeline are consistent with previous observations^15^: differences in transcriptional dynamics between different genes in human cells correlate with variation in bursting frequency (OFF times), while ON times distributions for different genes remain largely invariant. Overall, the results of these experiments indicate that HiTIPS can reliably measure the dynamics of transcription in an automated manner and for hundreds to thousands of live cells in long time-lapse experiments.

## Discussion

We developed HiTIPS as an open-source and automated image analysis software for large HTI datasets. Both individual researchers and imaging core facilities frequently struggle with identifying HTI software that combines ease-of-use for researchers with no programming experience with the flexibility to incorporate new analysis modules and algorithms for image bioinformatics developers. In developing HiTIPS, we strived to find a balance between these two needs. We believe that HiTIPS achieves this by taking advantage of a flexible GUI that uses the PyQT5 framework (Fig. 1), and by using a plugin software architecture that allows the incorporation of not only new algorithms for existing analysis modules (e.g. nuclear segmentation, spot finding, etc.), but also of completely new HTI analysis modules.

As proof of principle, we have used to HiTIPS for the analysis of HTI assays designed to address biological questions related to nuclear biology. We show that HiTIPS performs robustly for spot detection, fluorescence intensity, morphology, and kinetics measurements for nuclear compartments or markers at the single-cell level (Fig. 2). We used these HTI measurements to address a variety of questions related to 3D genome architecture (Fig. 3), centromere biology (Fig. 4), and transcriptional dynamics (Fig. 5). The results of these studies highlight HiTIPS ability to analyze data from HTI assays incorporating various fluorescent sample preparation techniques (FISH, IF, recombinant fluorescent proteins) and spanning multiple cellular processes.

We consider HiTIPS a valuable, user friendly addition to the suite of open-source software platforms available for HTI analysis ^32,33^. While HiTIPS does not currently include a vast range of analysis modules, this software introduces specific, custom algorithmic options for nuclei segmentation, nucleus tracking, spot detection, spot registration, spot tracking, and spot fluorescence intensity measurements. In addition, the modular nature of HiTIPS allows the future development of custom-tailored analysis tools for measurement of many biological features. In the future, we expect that novel, additional algorithms for FISH spot detection, such as FISH-quant ^17^, and for nuclear segmentation and tracking, could be added to HiTIF. Furthermore, as a natural progression for HiTIPS, we also foresee the addition of modules to segment the cell body and cell membranes, to extend the range of biological questions to other cellular compartments beside the nucleus. Similarly, we envision the potential addition of analysis modules in HiTIPS to measure fluorescence texture properties, additional morphological properties, and relational/cell neighborhood properties. Given the open-source and plugins software architecture of HiTIPS, we hope that the biological image analysis community will contribute to the development of these new HiTIPS features.

## Methods

### siRNA Oligos Transfection and Immunofluorescence

HCT116-Cas9 cells ^34^ were grown in RPMI-1640 medium (ATCC, Cat. No. 30-2001) supplemented with 10% fetal bovine serum (FBS, Gibco, Cat. No. 10-082-147) and maintained at 37°C in 5% CO2. siRNA Reverse transfection and Immunofluorescence staining were performed in a 384-well glass bottom plate (CellVis, Cat. No. P384-1.5H-N). The siRNAs were siNCAPH2 (Thermo Fisher Scientific, Cat. No. 4392420, Assay ID s26585), and siScrambled (Thermo Fisher Scientific, Cat. No. 4390847). For the reverse transfection, 150 nl of 5 mM siRNA and 50 nl of Lipofectamine RNAiMAX reagent (Invitrogen, Cat. No. 13778075) were individually diluted in 20 µl of serum-free OptiMEM medium (Thermo Fisher Scientific, Cat. No. 31985070) and sequentially added to each well. The siRNA and RNAiMAX mix was incubated for 30 min at RT. Cells were trypsinized, and a cell suspension (2000 cells in a volume of 20 µl) was prepared in OptiMEM supplemented with 20% FBS. 20 µl of the cell suspension was added to each well containing the RNAiMAX/siRNA oligo complexes. Transfected cells were grown for 72 hours in a cell incubator at 37°C and then fixed with 2% paraformaldehyde (PFA, Electron Microscopy Sciences, Cat. No. 15710) in PBS. Fixed cells were washed three times with PBS. Cells were then permeabilized using a 0.5% Triton X-100 (Milipore Sigma, Cat No. 9036-19-5) solution in PBS for 15 minutes at RT, washed three times with 50 µl PBS, and blocked in a 5% BSA (Milipore Sigma, Cat No. 9048-46-8) solution in PBS for 15 minutes at RT. Immunofluorescence staining against the centromeric protein CENPC was performed using a primary CENPC antibody (MBL Bio Science, Cat. No. PD030, raised in Guinea pig) at 1:1000 dilution for 1 hour at RT, and a Goat Anti-Guinea pig IgG H&L secondary antibody (AlexaFluor 488, Abcam, Cat. No. Ab150185) at 1:500 dilution for 1 hour at RT. For nuclear staining, 40 µl of a 5 mg/ml DAPI 4′,6-diamidino-2-phenylindole (DAPI, Thermo Fisher Scientific, Cat. No. 62248) solution in PBS were added to each well.

### High-Throughput DNA FISH

HCT116 RAD21-mAID-mClover (RAD21-mAC) cells ^25^ were cultured at 37 °C in 5% CO2 in McCoy’s 5A medium supplemented with 10% FBS, 2 mM L-glutamine, 100 U/ml penicillin, and 100 µg/ml streptomycin. For the FISH experiment, cells were plated at a density of 8000 cells per well in 384-well imaging plates (PhenoPlate 384-well, Revvity, Cat. No. 6057500) and allowed to grow overnight. The following day, the medium was replaced with either supplemented medium containing 170 mM Auxin (Sigma-Aldrich, Cat. No. I3750) to induce the degradation of RAD21, or with medium with an equivalent amount of DMSO alone as vehicle control. The cells were then incubated with or without Auxin for 3 hours and fixed in 4% PFA (Electron Microscopy Sciences, Cat. No. 15710) in PBS for 10 minutes. After fixation, the plates were rinsed three times in PBS and stored in PBS at 4 °C.

We conducted high-throughput fluorescence in situ hybridization (hiFISH) as previously described ^4,35^. BAC FISH probes were selected to hybridize to the boundary regions of the topologically associated domain (TAD) on chromosome 7 containing the *EGFR* gene. Fluorescently labeled BAC probes were generated by nick translation at 14°C for 1 hour and 20 minutes. The reaction mixture included 40 ng/ml DNA, 0.05 M Tris-HCl pH 8.0, 5 mM MgCl2, 0.05 mg/ml BSA, 0.05 mM dNTPs (including fluorescently tagged dUTP), 1 mM β-mercaptoethanol, 0.5 U/ml E. *coli* DNA Polymerase, and 0.5 mg/ml DNase I. The reaction was stopped by adding 1 µl of EDTA per 50 µl reaction volume, followed by a heat shock at 72 °C for 10 minutes. Probes were labeled either with DY549P1-dUTP (Dyomics, Cat. # 549P1-34) or with DY647P1-dUTP (Dyomics, Cat. # 647P1-34). The reaction was then stored at −20 °C overnight. Next, the two probes (0.5 mg per probe) were combined, precipitated with ethanol, and resuspended in 14 µl of hybridization buffer (50% formamide pH 7.0, 10% Dextran Sulfate, and 1% Tween-20 in 2X SSC) per well. Cells were rinsed twice with PBS and subjected to permeabilization. Permeabilization was performed at room temperature for 20 min using 0.5% w/v saponin/0.5% v/v Triton X-100 in PBS. After rinsing the cells with PBS twice, cells were deproteinated for 15 minutes in 0.1 N HCl and neutralized for 5 minutes in 2X SSC at room temperature. Cells were equilibrated overnight in 50% formamide/2X SSC at 4°C. The probe mix was warmed to 72 °C prior to the hybridization reaction. Next, 14 µl/well of resuspended probe mix was added to the plate and denatured at 85 °C for 7 min, followed by immediate transfer to a 37 °C incubator for a 48-hour hybridization period. Post-hybridization, plates were rinsed once at room temperature with 2X SSC, followed by three rinses with 1X SSC and 0.1X SSC, all warmed to 45 °C. Cells were stained with 3 µg/ml DAPI for 15 minutes, then rinsed and mounted in PBS and subsequently imaged on a high-throughput confocal microscope.

### High-Throughput Live Cell Imaging of Transcription

For the live cell transcription assay, human bronchial epithelial cell lines (HBEC3-KT) with a monoallelic insertion of an MS2 array in the intron of the model genes were used, as previously described ^15^. To enable visualization of the nascent RNA, the viral MS2 capsid protein (MCP) fused to GFP, and an NLS (nuclear localization signal) tag were stably introduced into the cells using lentiviral expression vectors ^36^. Cells were grown in Keratinocyte serum-free medium (Thermo Fisher Scientific, Cat. No. 17005042) supplemented with bovine pituitary extract (Thermo Fisher Scientific, Cat. No. 13028014) and human growth hormone (Thermo Fisher Scientific, Cat. No. 1045013). For imaging experiments, cells were cultured in 384 well plates (PhenoPlate 384-well, Revvity, Cat. No. 6057500) and imaged at 37°C, 5% CO2, 80% humidity.

### High-Throughput Image Acquisition

High-throughput imaging was performed using either a Yokogawa CV7000 or a CV8000 high-throughput spinning disk confocal microscopes.

For DNA FISH experiments, we used 405 nm (DAPI Channel), 561 nm (Probe A channel), or 640 nm (Probe B channel) excitation lasers. In addition, we used a 405/488/561/640 nm excitation dichroic mirror, a 60X water objective (NA 1.2), and 445/45 nm (DAPI Channel), 600/37 nm (Probe A channel), or 676/29 nm (Probe B channel) bandpass emission mirrors in front of a 16-bit sCMOS camera (2048 × 2048 pixels, binning 1×1, pixel size: 0.108 microns). Z-stacks of 7 microns were acquired at 1 micron intervals and maximally projected on the fly. Images were acquired in 32 fields of view (FOV) per well.

For IF experiments, we used 405 nm (DAPI Channel) or 488 nm (CENPC channel) excitation lasers, a 405/488/561/640 nm excitation dichroic mirror, a 60X water objective (NA 1.2), 445/45 (DAPI Channel) or 525/50 nm (CENPC Channel) bandpass emission mirrors, and a 16-bit sCMOS camera (2048 × 2048 pixels, binning 1×1, pixel size: 0.108 microns). Z-stacks of 14 microns were acquired at 1 micron intervals and maximally projected on the fly. Images were acquired in 22 FOV per well.

For live cell imaging experiments, we used a 488 nm excitation laser, a 405/488/561/640 nm excitation dichroic mirror, either a 40X air objective (NA 0.95) or a 40X water objective (NA 1.15), a 525/50 nm bandpass emission mirror, and a 16-bit sCMOS camera (2048 × 2048 pixels, binning 2×2, pixel size: 0.325 microns). Z-stacks of 0.5 microns were acquired at 100 sec intervals and maximally projected on the fly, every 100s for 10 hours.

In all cases, images were corrected on the fly with Yokogawa proprietary software to subtract the camera dark background, and to compensate for illumination artifacts (vignetting), and for chromatic aberrations and cameras alignment.

### HiTIPS Implementation

HiTIPS uses the PyQt5 Python module, which offers a user-friendly GUI enabling interactive data analysis. Its architecture implements multiprocessing to optimize computational efficiency, a critical aspect when dealing with large-scale bioinformatics datasets. The parallel processing scheme in HiTIPS is designed to completely analyze (Nuclei segmentation, spot detection etc.) each FOV in a separate thread. Depending on the available hardware, parallel processing in HiTIPS can reduce the analysis time by 5-to 8-folds, depending on the analysis workflow.

HiTIPS depends on several Python scientific computing libraries, including numerical computation and data manipulation (Scipy, pandas), image processing (Pillow, Matplotlib, imageio, scikit-image, and OpenCV), dynamic cell tracking (btrack), machine learning-based image segmentation and classification (DeepCell ^37^ and CellPose ^38^), image input/output and format conversion (aicsimageio, nd2reader), and Hidden Markov Model fitting (hmmlearn). At least 8 GB of RAM are required to run HiTIPS, but having 32 GB of RAM may be required for larger FOVs and for 3D volumes. In addition, when using deep learning based nuclear segmentation or cell tracking models in HiTIPS (i.e., CellPose and DeepCell), the availability of graphical processing units (GPUs) substantially improves the inference speed of these models.

### Nucleus Segmentation

Nucleus segmentation using images of nuclei stained with a fluorescent dye or a recombinant fluorescent nuclear protein is the key first step in the vast majority of HTI analysis pipelines. Given the high relevance of this step for HTI, a substantial amount of work in the field has been devoted to making nuclear segmentation algorithms fast, precise, and robust to fluctuations in cell confluency and to heterogeneity in nucleus morphology across different cells ^39^. For this reason, and to take advantage of previous advances made by other groups, we focused on integrating existing state-of-the-art nucleus segmentation algorithms into HiTIPS so that end users can easily access them and modify their parameters if needed. Accordingly, the HiTIPS GUI allows users to choose among a traditional CPU based method (Algorithm 1) for segmentation in the nucleus segmentation module, and two recent deep –learning-based methods, CellPose ^40,41^ and DeepCell ^42^. Deep learning-based nuclei segmentation models do not involve time consuming parameter optimization, and they generally provide excellent segmentation performance on a variety of different cell lines “out-of-the-box”. On the other hand, the speed performance of segmentation models really benefits from access to graphical processing units (GPUs), which tend to be expensive and difficult to setup for end users. Traditional image processing algorithms for nucleus segmentation can be fast if properly optimized and can handle a variety of edge cases upon expert parameter optimization. The watershed-based segmentation method is the CPU-based approach integrated into HiTIPS. It starts with image padding and noise reduction via median filtering, followed by image binarization using Li’s iterative method ^43^. The binary image is then processed using morphological operations and a Gaussian kernel to connect fragmented nuclei. The method labels connected components and it calculates the center of mass for each, creating a new mask image. A watershed transform ^44^ is applied using this mask and the distance-transformed image to separate adjacent nuclei effectively. Finally, a boundary image is created, resized to the original size, and any holes are filled to generate the final mask. By providing an easy selection of different nuclear segmentation methods via a GUI, HiTIPS allows users to choose and optimize the method that works best on their images, and in the context of the available computational hardware infrastructure.

### Spot Detection

HiTIPS includes morphologic, intensity, and filtering-based approaches for fluorescent spot detection. Currently, HiTIPS incorporates four different spot detection methods: Direct Thresholding, Gaussian Filter, Gaussian Laplacian, and Enhanced Gaussian Filter and Laplacian. The spot detection methods offered as part of HiTIPS provide have their own set of strengths and limitations, which need to be considered when choosing the appropriate spot detection method for a given type of biological sample and imaging assay.

The Direct Thresholding method applies a direct thresholding technique for spot segmentation without any filtering. It is a straightforward and computationally efficient approach suitable for scenarios where spots have large contrast. However, this method may be less effective when dealing with spots that have low contrast or are close to the background intensity level. Additionally, it has limited capability to handle spots with varying intensity gradients. The Gaussian Filter method utilizes a Gaussian filter to reduce noise and enhance spots. This method performs well when spots have a relatively uniform intensity distribution and works better when spots are close together or overlapping compared to the Gaussian Laplacian method. However, it may be less effective in enhancing spots with sharp intensity variations or irregular shapes. Careful consideration of parameters such as the Gaussian filter size (sigma) and thresholding parameters is necessary. The Gaussian Laplacian method enhances spots by applying a Gaussian Laplacian filter to the input image and then segments the spots using thresholding. By utilizing the negative lobes of the Gaussian Laplacian kernel, this method not only enhances the spots but also removes the background around the spot, improving the effectiveness of automatic thresholding. It is a relatively simple and computationally efficient method. However, it may face challenges when spots are closely located or overlapping due to limited resolution. Sensitivity to parameters such as the Laplacian filter size (sigma) and thresholding parameters should be considered. The Enhanced Gaussian Filter and Laplacian method, combines the strengths of both the Gaussian Filter and Gaussian Laplacian methods. It first applies a Gaussian filter to the input image, followed by a Gaussian Laplacian filter on the filtered image, and it finally uses fluorescence thresholding for spot segmentation. This method provides enhanced capabilities for detecting spots with varying intensity gradients and can improve overall spot detection accuracy. However, achieving optimal results may require careful tuning of filter sizes (sigma), and the choice of thresholding parameters may still impact its performance.

The spot detection methods provided in HiTIPS enable the detection and localization in the X and Y dimensions of fluorescent spots generated by DNA/RNA FISH staining, or from other biological structures in maximally projected 3D z-stacks microscopy images. Subsequently, maximum intensity or Gaussian-fitted maximum intensity can be employed to estimate the spot center positions in the Z dimension of the z-stack.

### Nuclei Tracking

Nuclei tracking can be framed as a linear assignment problem in which N_i_ objects in frame i are matched up with N_i+1_ objects in frame i+1. Shadow objects can be introduced to account for births (i.e. from cell division events) or deaths (i.e. cells leaving the field of view). We incorporated two cell tracking methods in HiTIPS to accommodate HTI assays using cell lines with different levels of confluency and mobility.

The first method we adopted ^45^ revisits and updates the Kalman filtering algorithm ^46^ and uses a Bayesian framework to improve the cell tracking accuracy and reliability. At the onset, the algorithm constructs tracklets, which are links between consecutive cell detections that do not exhibit cell division events. These tracklets from a prior frame are paired with observed cells in the current frame to form a Bayesian belief matrix, which initially holds a uniform probability of associations. Crucially, each tracklet deploys its own Kalman filter to predict the future state of a cell, basing its predictions on motion models and information from a cell state classifier. This classifier discerns nuclear morphological variations and chromatin condensation levels, which are crucial visual features in tracking. Belief updates in the matrix consider both motion evidence (using a constant velocity model) and appearance evidence (through a cell state transition matrix). The motion aspect focuses on the estimated future position of a cell, while the appearance aspect evaluates linking probabilities based on the state transitions determined by the classifier. Notably, this combination method aids in accurately identifying instances like cell divisions. After forming these tracklets, a global optimization approach employing multiple hypothesis testing is used. It constructs a large number of hypotheses based on the appearance and motion features, with the aim of identifying the most optimal track-linking solution. Hypotheses account for diverse cell behaviors, including cell divisions, false-positive tracks, or apoptosis events. The optimal set of hypotheses is determined using a maximization function, resulting in the amalgamation of tracklets into final tracks. Ultimately, a graph-based approach is leveraged to assemble these tracks into lineage trees, outputting a set of additional measurements such as generational depth and cell lineage.

The second tracking method adopted by HiTIPS is DeepCell ^42,47^. DeepCell employs a fully connected neural network that considers various features of each cell, including its appearance, local neighborhood, morphology, and motion. These pieces of information are fed into the neural network, which then processes and summarizes them into a vector representation using a deep learning sub-model. To determine the relationship between cells in consecutive frames, the information from the past frame and the current frame is utilized. The Hungarian algorithm ^48^, is employed for this purpose, which is a combinatorial optimization algorithm that assigns the best possible associations between cells across frames. It determines whether the current cell is the same as a cell in the previous frame, a different cell, or a child cell derived from the cell in the previous frame. By combining the DeepCell neural network with the Hungarian algorithm, this tracking method aims to accurately track and link cells across frames, considering their various characteristics and relationships.

### Nuclei/RNA Spot Image Patch Generation

Each cell track, representing the same nucleus monitored across the time-lapse movie, is precisely cropped from the full FOV into 128 × 128 pixels image patches. At each time point, the cropping algorithm positions the center of mass of the segmented nucleus ROI at the center of the cropped image, thus optimizing the positioning and the orientation of the nuclei, which is further refined in the subsequent nucleus registration step. While it is possible to segment and track partial nuclei ROIs from the moment they enter the full FOV, until they partially exit it, this approach carries the risk of missing crucial objects or events in the nucleus. This can also potentially decrease the precision of frame-to-frame nuclei registration. Accordingly, HiTIPS provides the option to select only nuclei which are entirely within the frame, thereby improving the accuracy of tracking and RNA spot detection.

### Nuclei/RNA Spots Image Patch Registration

Accurate frame-to-frame rigid and rotational registration of the nucleus ROI is indispensable to track the dynamics of spots signals in the same nucleus over time. For example, in the time-lapse images of MS2/MCP-GFP labelled transcription sites, nuclei segmentation is often performed using the nucleoplasmic fluorescence background of the NLS-tagged MCP-GFP protein, which contains very little to no information about other sub-nuclear structures, such as nucleoli or chromocenters. As a consequence, feature-based image registration methods such as SIFT or SURF ^49–51^, which rely on the presence of prominent texture features that remain consistent over time, cannot be utilized. To overcome this limitation, HiTIPS incorporates two novel registration methods that correct for nucleus translation and rotation across time-lapse movies. This is achieved by taking into account subtle variations in nuclear shape across the cropped ROI time-lapse movie in a two-step process (Algorithm 2 and Algorithm 3). These methods provide an effective solution for tracking nuclei positioning across frames, thus facilitating the successful tracking of transcription sites in live cells.

The first method (Algorithm 2) starts with setting the angle between the major axis of the nucleus (α) and the horizon as zero. For each iteration greater than zero, two variables, α’ and α”, are initialized at 0 and 180 degrees, respectively. The algorithm then computes eight specific features, namely, Cosine Similarity Index, Mutual Information, Structural Similarity Index, Mean Square Error, Variation of Information, Adaptive Random Error, and Peak Signal to Noise Ratio for these initial values of α’ and α”. If the majority of the features for α’ exceed those for α”, then the value of α for this iteration is set as α’, else it is set as α”. This is followed by a process where α’ and α” are set to α ± i (where i varies from 1 to 5) and a similar comparison of features is performed to update α. The algorithm then increments the iteration count (n), and repeats these steps until the final step, where the spot coordinates are mapped to the nuclei patch and used for spot tracking.

The second method (Algorithm 3) begins by assigning an initial rotation of 15 degrees to α to prevent cumulative drift. Next, a series of transformations is performed on the centered nuclei including median filtering, upsampling, and polar warping. Following these transformations, subpixel image translation registration is carried out by cross-correlation in the polar Fourier domain. The algorithm then corrects for the initial rotation assigned in step one by subtracting it. The final step involves mapping the spot coordinates to the nuclei patch and tracking these spots using Algorithms 5 and 6.

The intensity-based registration approach (Algorithm 2) has been proven to work better when cells shape changes during along the movement on their trajectory, however, large frame-to-frame intensity variations can introduce angle shift or translation in the registration results. On the other hand, the Fourier phase transform-based approach (Algorithm 3) is more robust to intensity change and less robust to frame-to-frame shape deformation.

### Assignment of Transcription Spots to Timelapse Tracks

The assignment of individual fluorescent spots detected at different time points to common tracks using hierarchical clustering integrates several algorithms for optimal results. HiTIPS employs a two-step process to effectively identify and organize spatial clusters of spots within each cell in projected stacks of images across the time dimension. The first step (Algorithm 4) calculates the pairwise Euclidean distance between all transcription spots in the time-projected image, and then uses single-linkage hierarchical clustering to generate an initial set of labels for the clusters. This algorithm also identifies the centroids of each unique cluster label, it calculates the standard deviation of distances from the centroid for each cluster, and it identifies outlier spots that exceed a user-defined threshold distance from the centroid of the clusters. Once the outliers are identified they are arbitrarily labelled with a label of “zero”. The second algorithm (Algorithm 5) first determines the size of each cluster (i.e. the number of spots in the cluster) and separates them into “large” and “small” categories based on a user-defined threshold. Then, Algorithm 5 calculates the Euclidean distances between points for each small cluster and all points in all the other large clusters. If the distance between the closest point in a large cluster is less than a user defined distance, the label of that closest point is assigned to the points in the small cluster. If not, it is labeled as an outlier. Through this process, small clusters and outliers are effectively merged into larger, more significant clusters, which streamlines the data structure and improves the interpretability of the results. The updated cluster labels, which correspond to tracks, for example of MS2/MCP-GFP representing sites of active gene transcription, across time, are then returned.

### Integrated intensity measurement

The integrated fluorescence intensity measurement of spots includes two components: Local background estimation and Gaussian mask fitting ^52^. The Local Background Estimation Algorithm (Algorithm 6) first addresses the preprocessing of the images. This algorithm utilizes a least squares method to fit a background 2D plane using the fluorescence intensity values of the pixels at the border of an 11 × 11 pixels matrix centered around the location of the spot. The estimated background plane is then subtracted from the original image, thus locally correcting for potential non-uniform illumination, and compensating for systematic imaging noise.

Following the background correction, the Gaussian Mask Fitting Algorithm (Algorithm 7) fits a Gaussian mask to the image to isolate and analyze individual spots within the image. The Gaussian mask can be either statically applied based on a given centroid or it can be iteratively adjusted to improve the accuracy of the fitting. The process involves iterative computation and adjustment of the centroid coordinates of the Gaussian mask until the difference between the old and new centroids becomes negligible, or until the maximum iteration count is reached. The final output from this process is the centroid coordinates of the Gaussian mask and the estimated photon number, which can then be used for further intensity track analysis.

### Image Processing Algorithms

#### Algorithm 1

CPU-Based Nuclei segmentation.

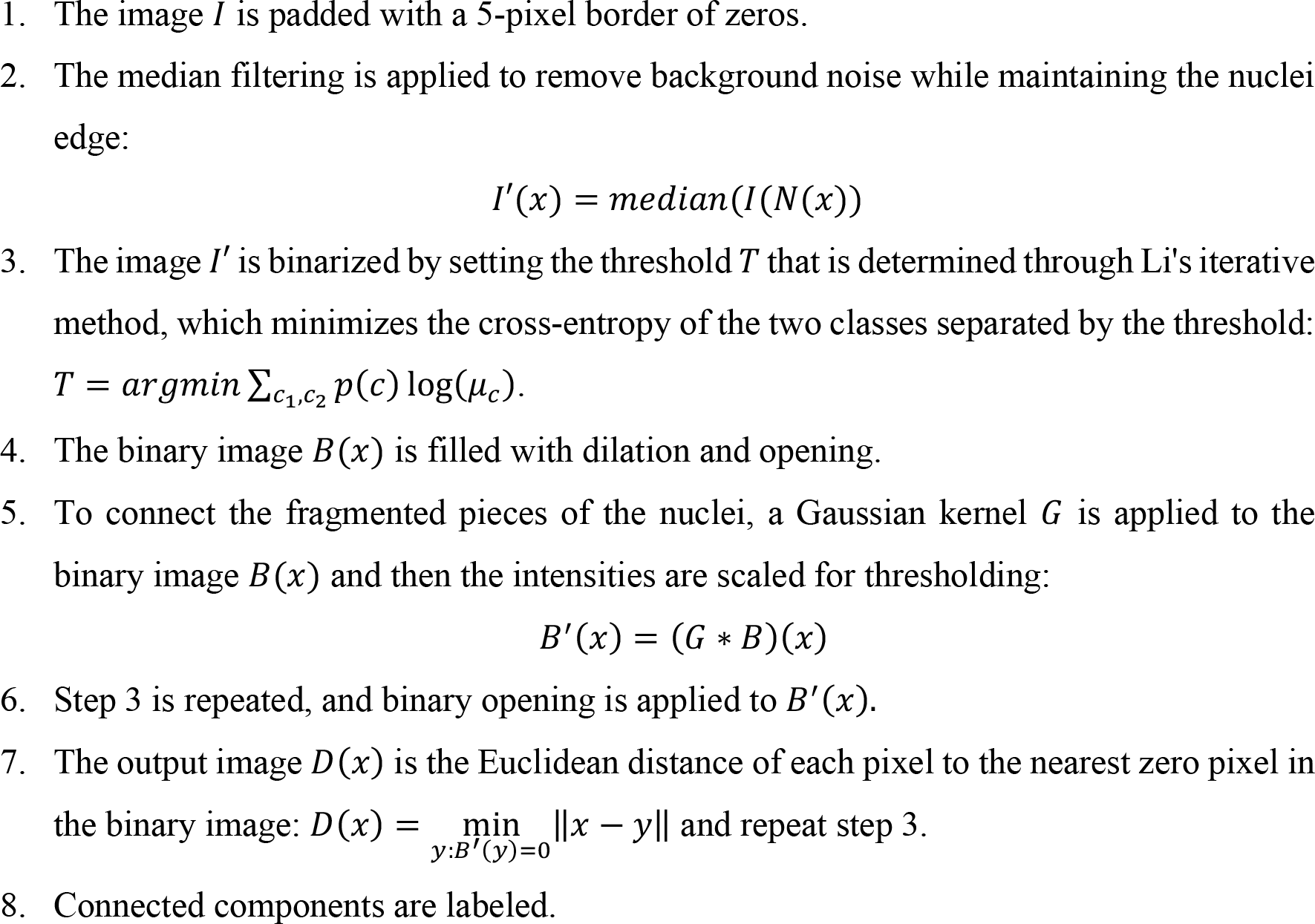
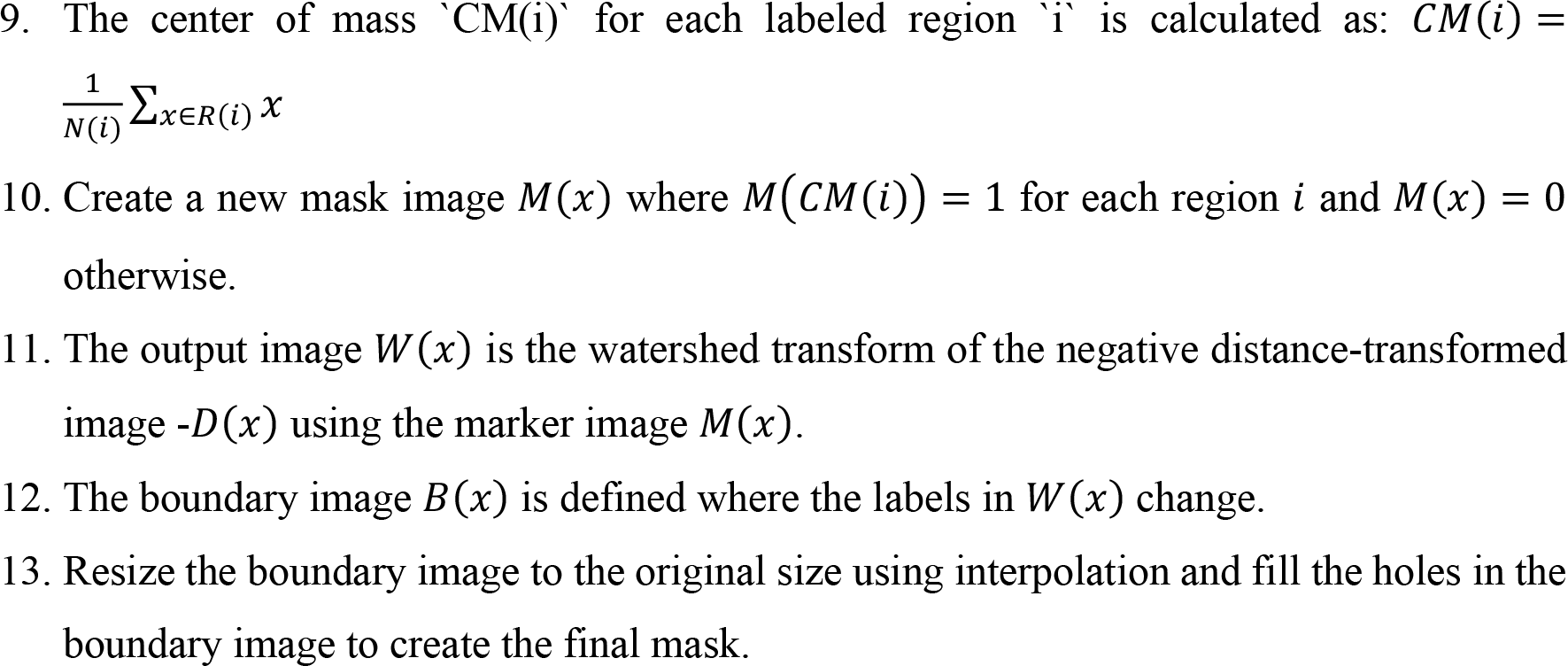

#### Algorithm 2

Intensity based timelapse nuclei alignment and RNA/gene assignment.

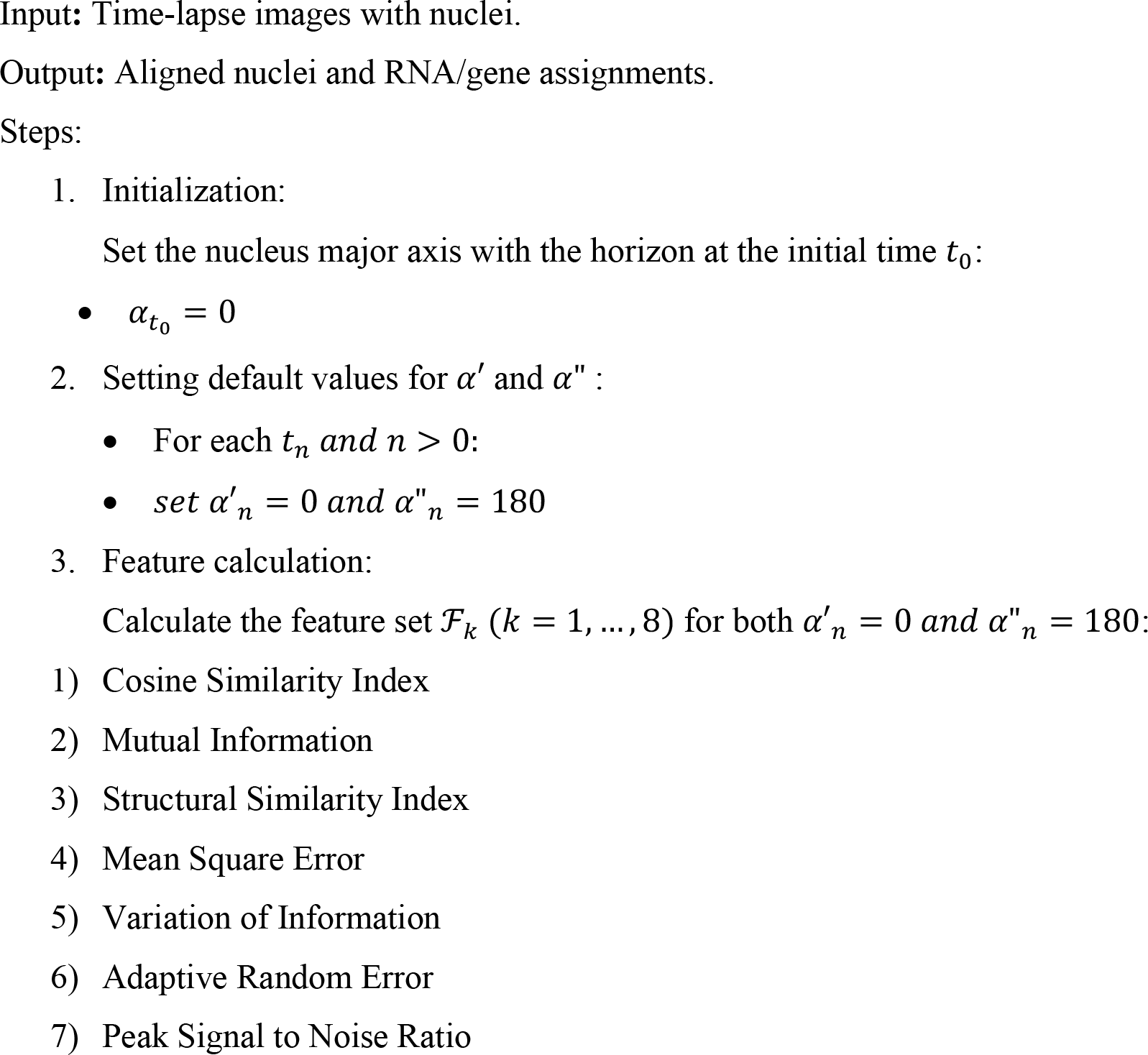
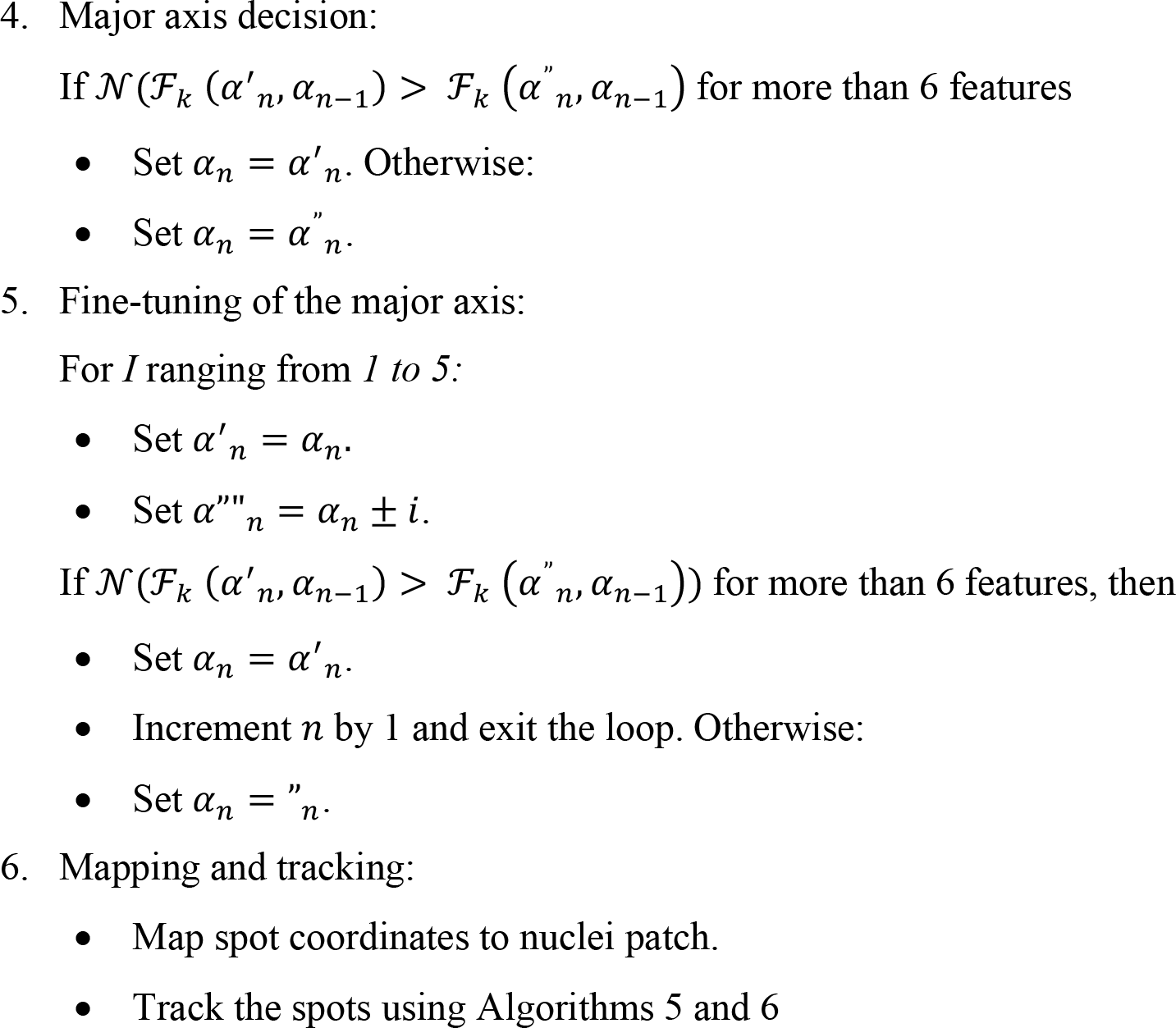

#### Algorithm 3

Phase-transform timelapse nuclei alignment and RNA/Gene assignment.

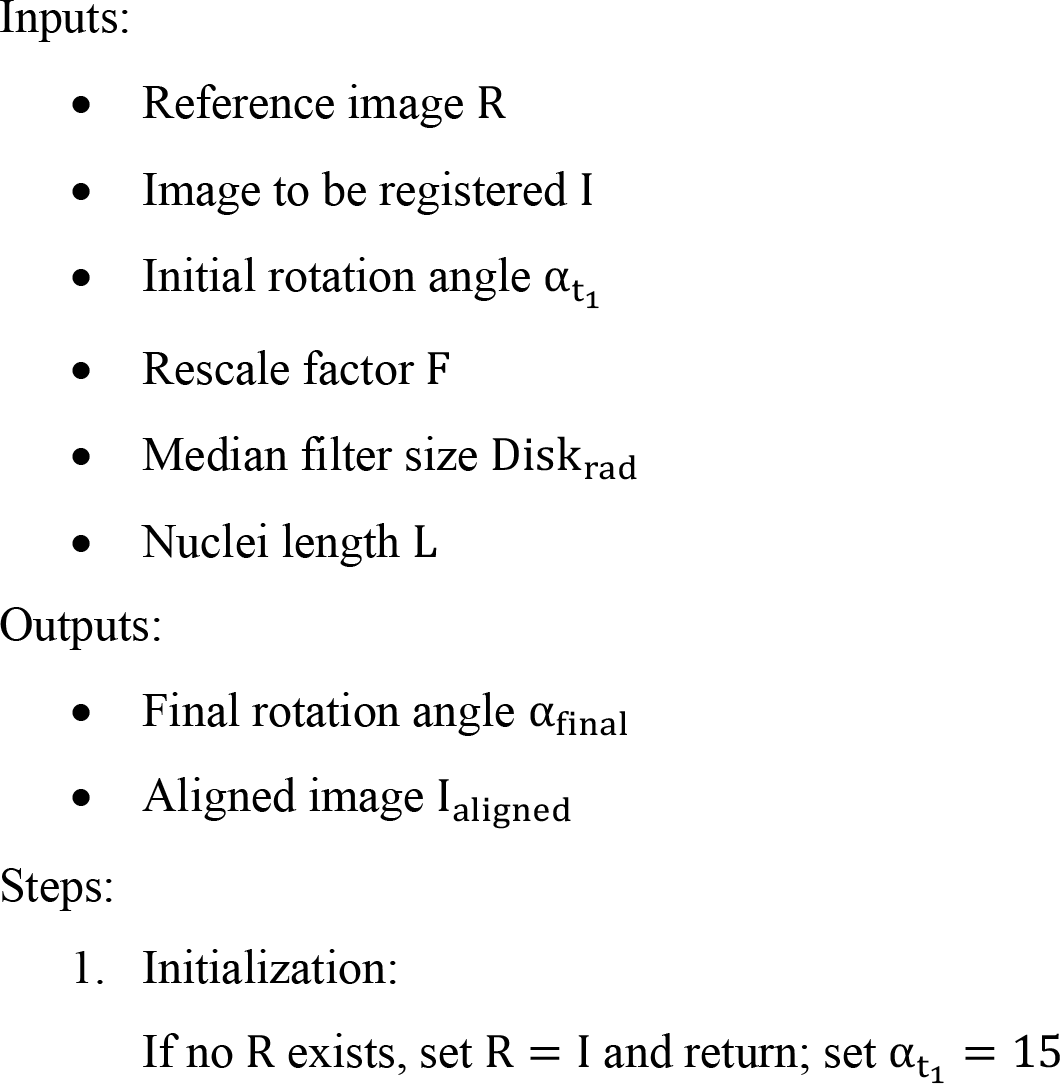
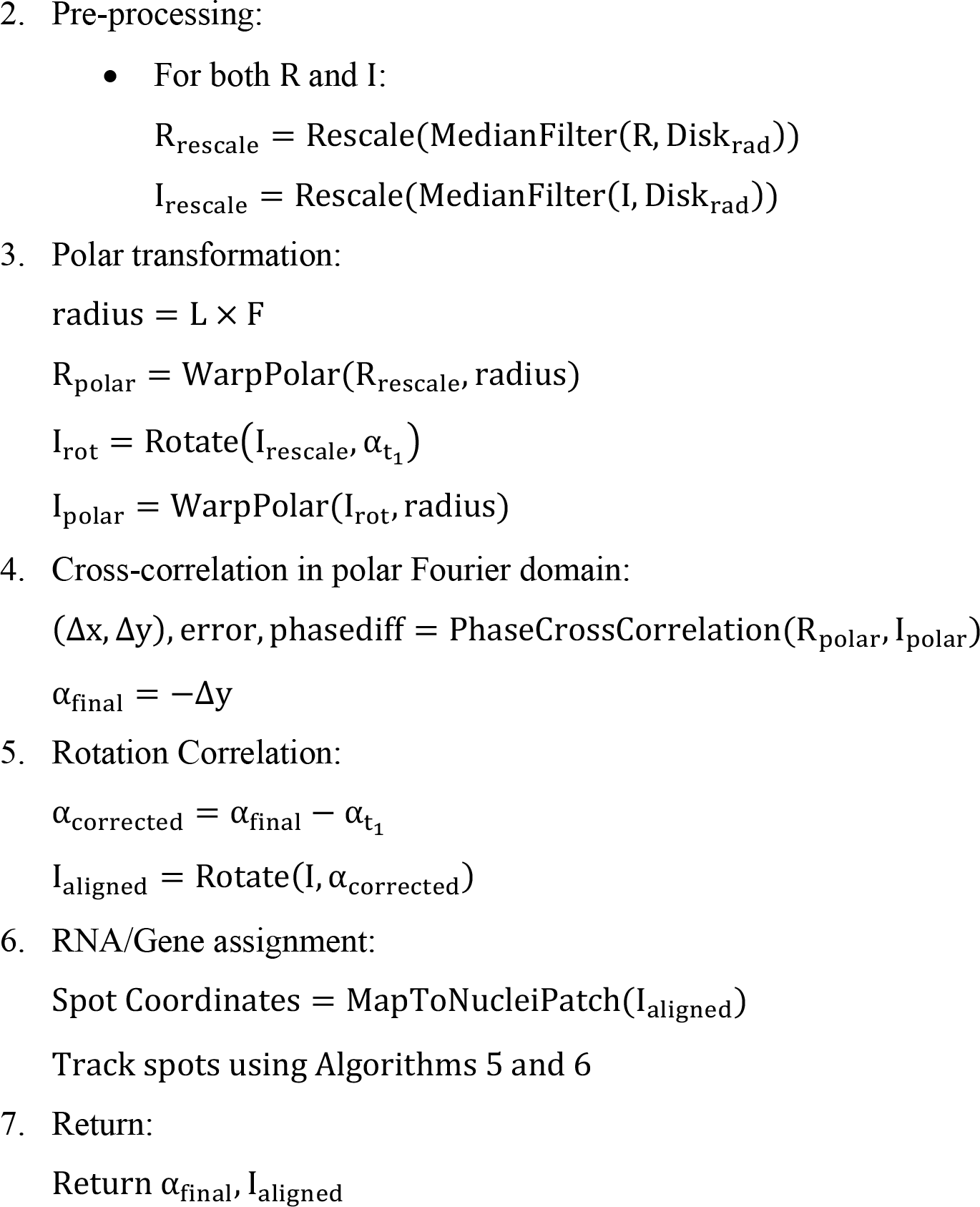

#### Algorithm 4

Hierarchical clustering and outlier removal algorithm.

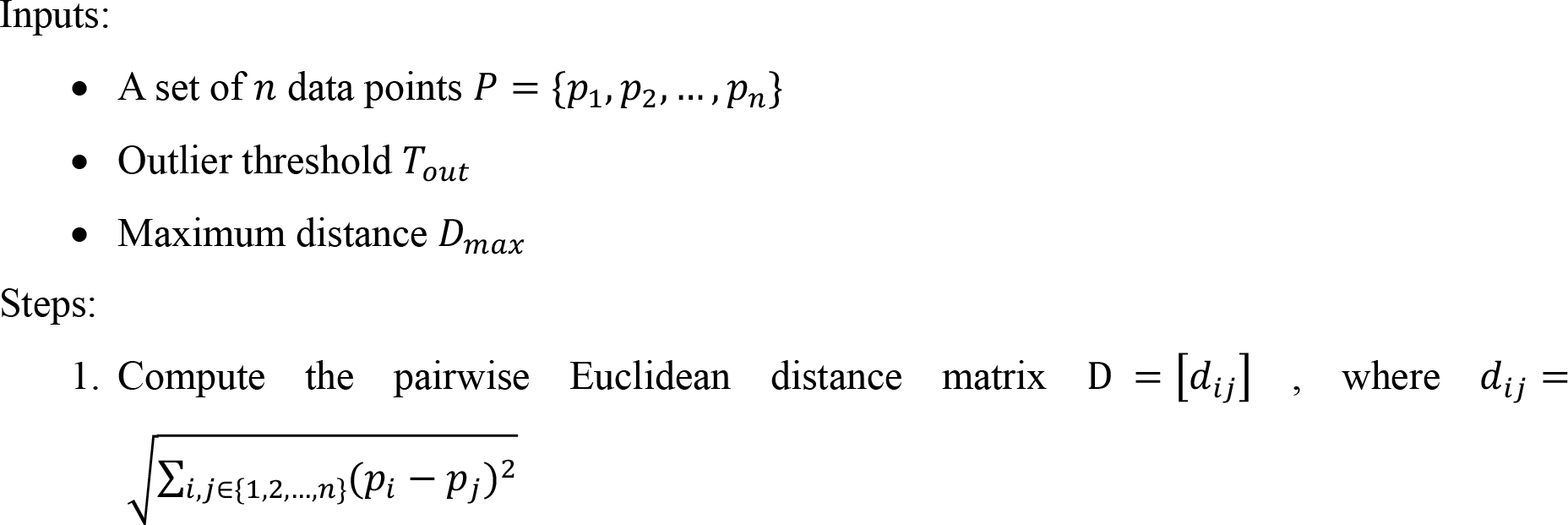
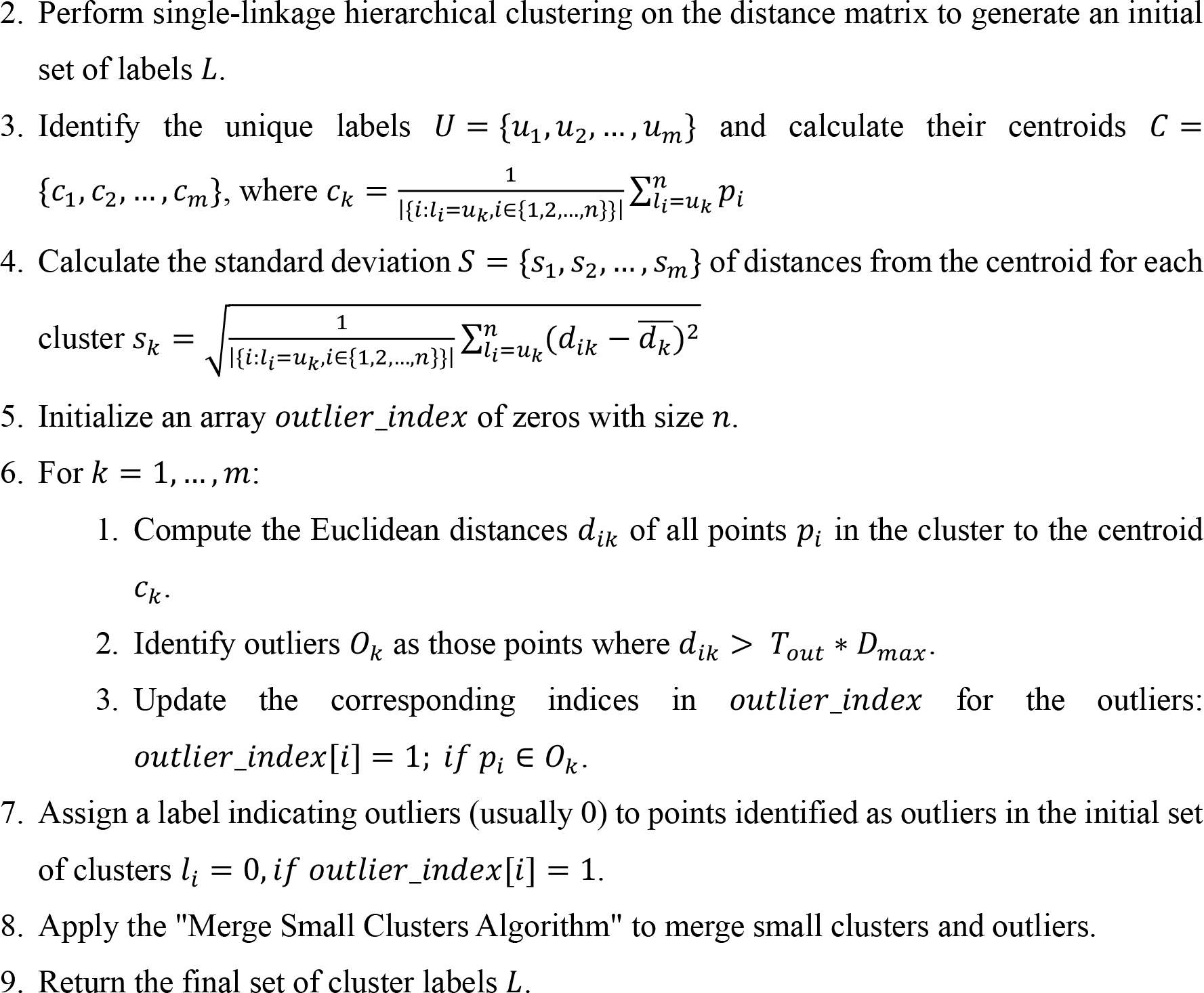

#### Algorithm 5

Merge small clusters algorithm.

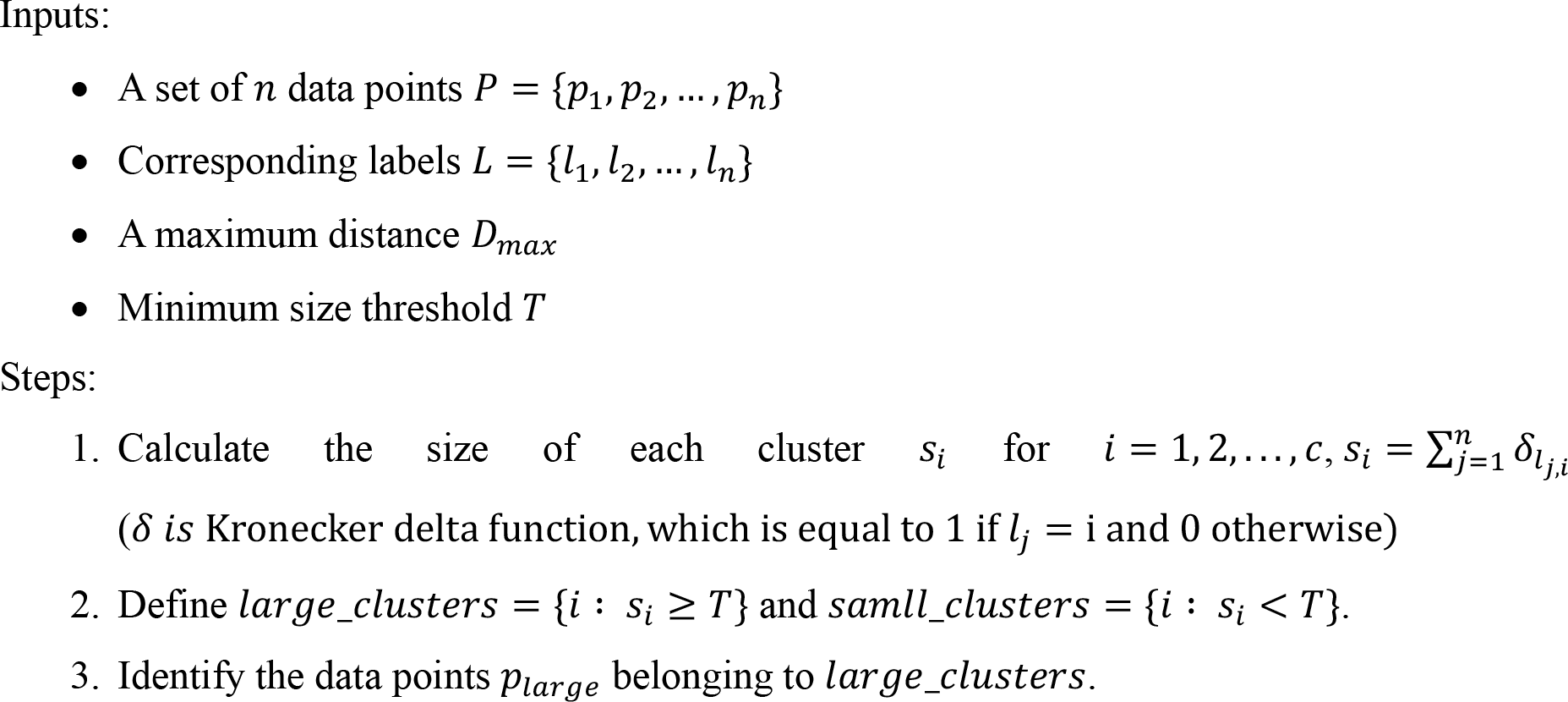
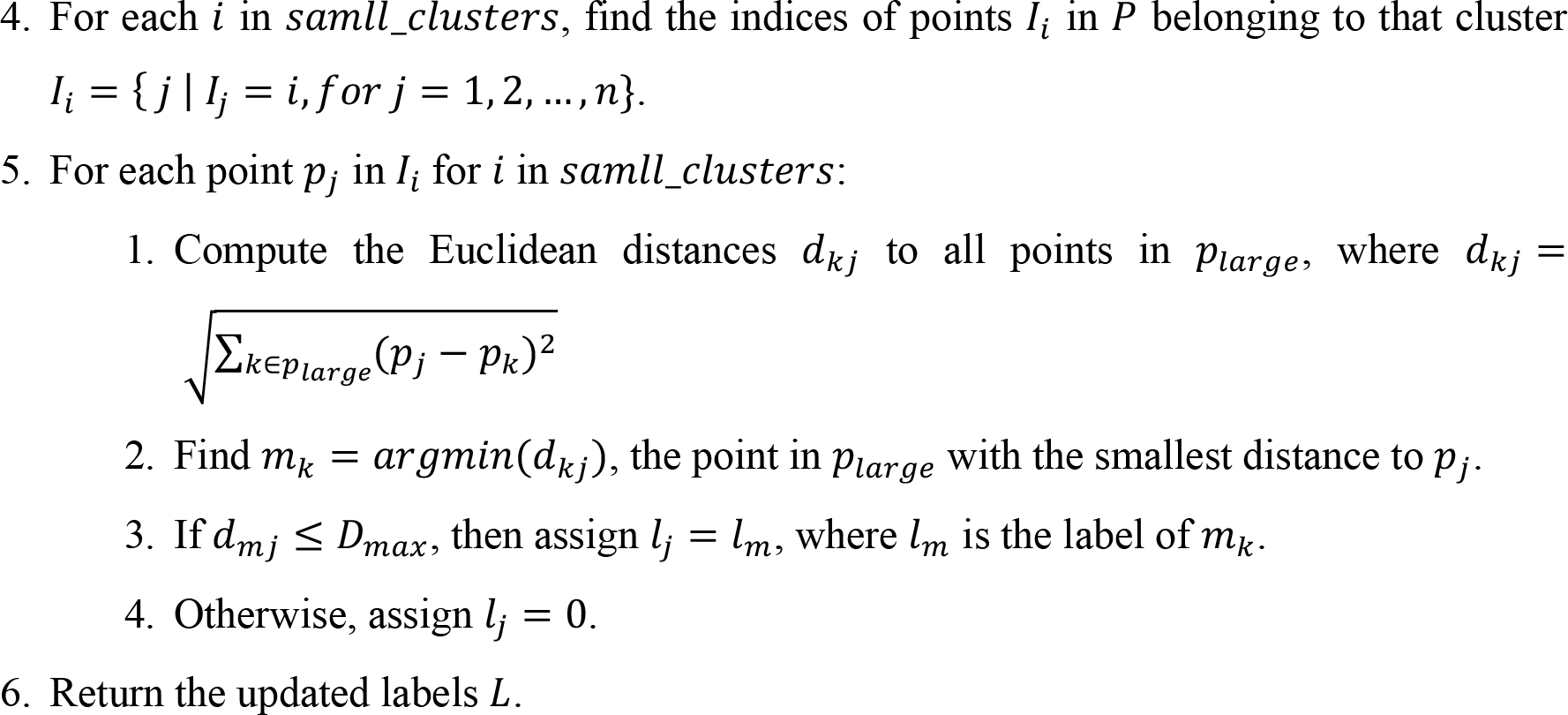

#### Algorithm 6

Local background estimation algorithm

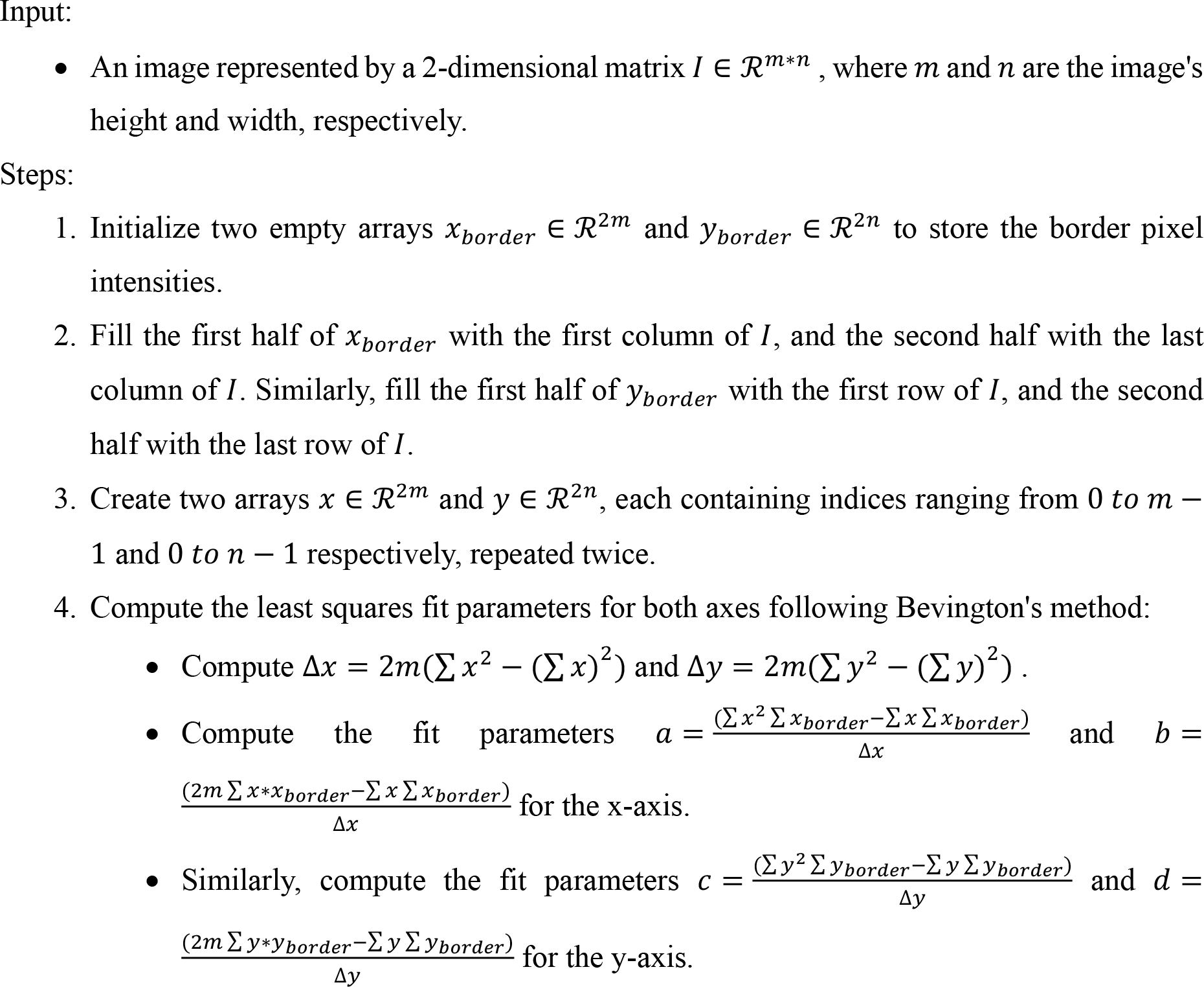
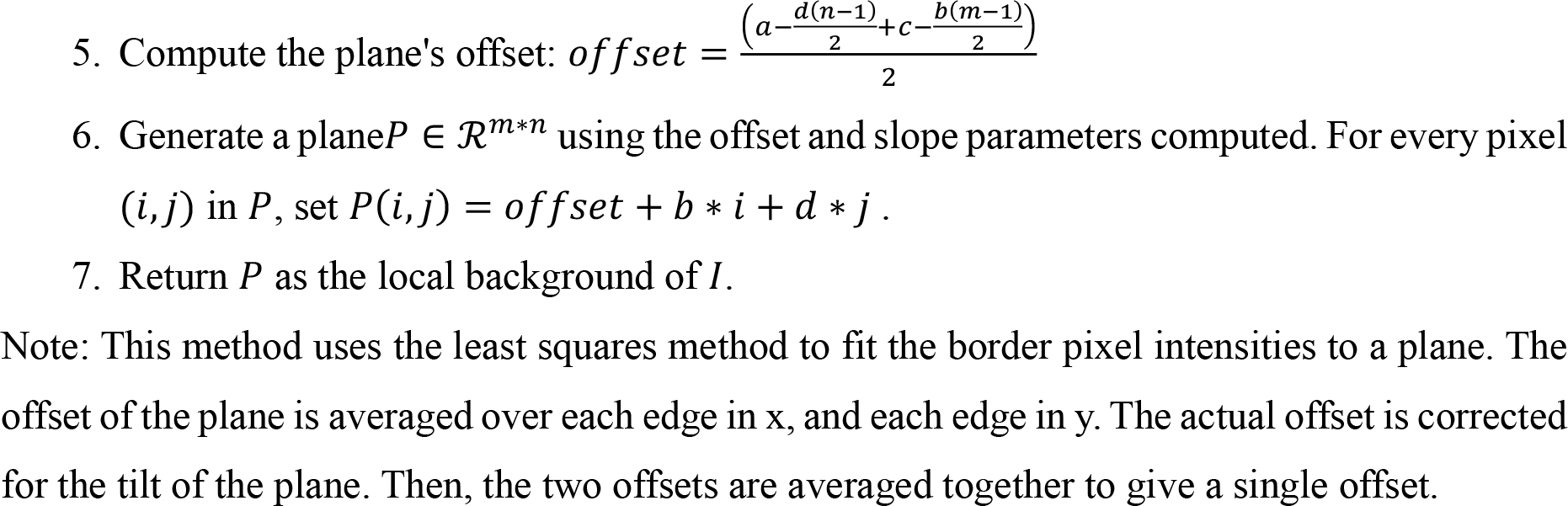

#### Algorithm 7

Gaussian mask fitting algorithm.

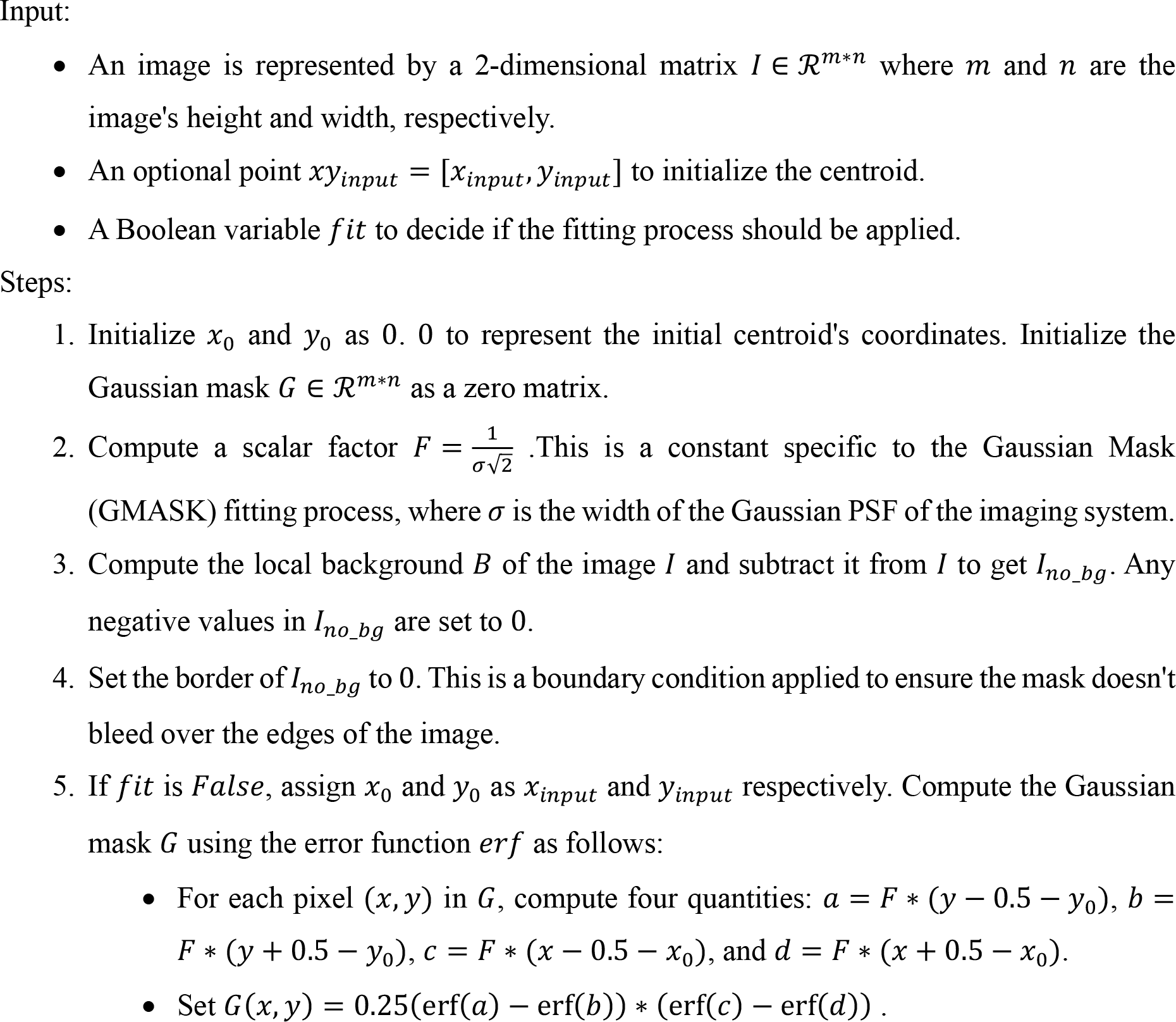
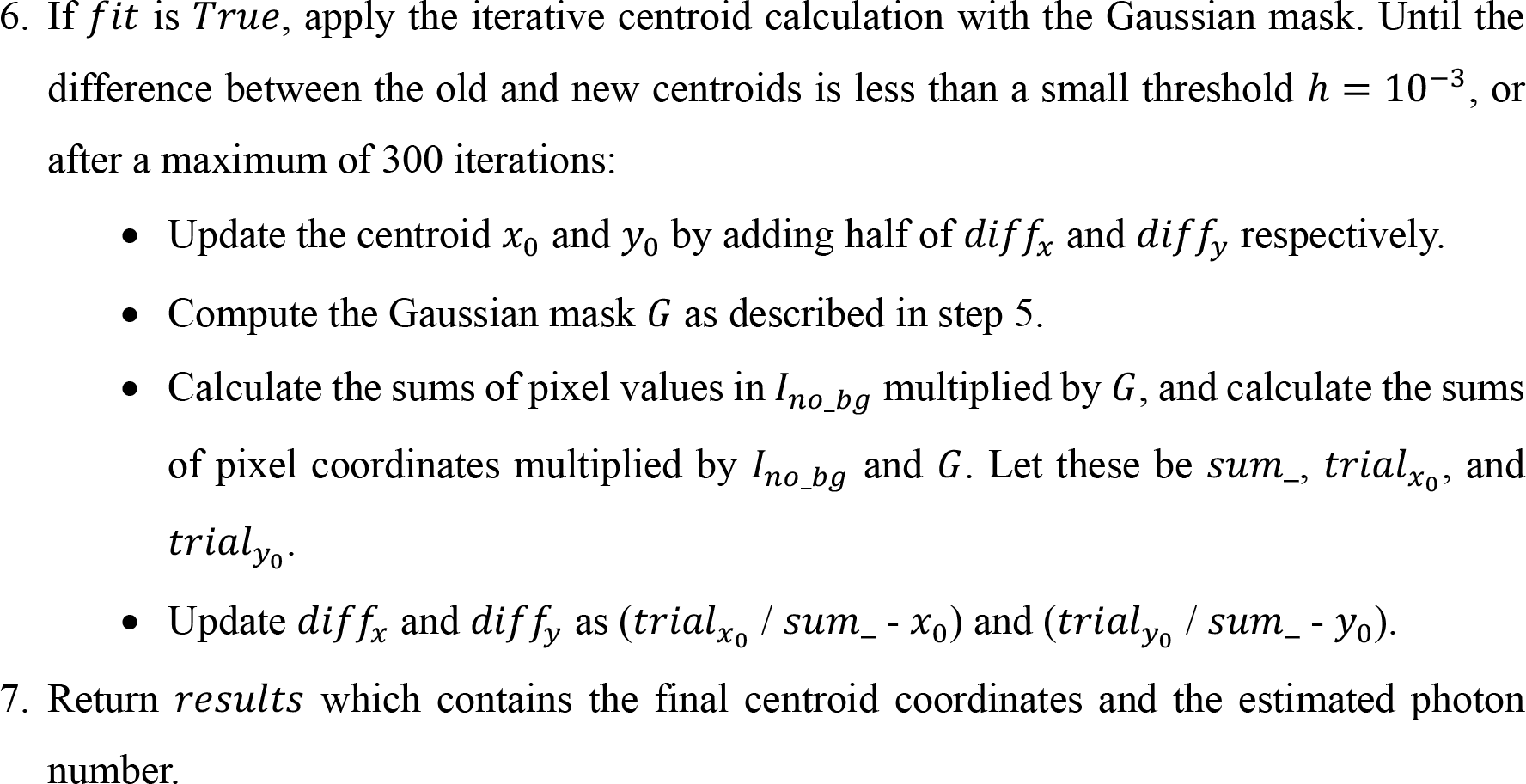

### CENPC Clustering Score Calculation

In our analysis, we employ a derivative of Ripley’s K-function ^30^, specifically designed to estimate the degree of spatial clustering at the single-cell level. This statistical measure, denoted as K(r), is defined as:

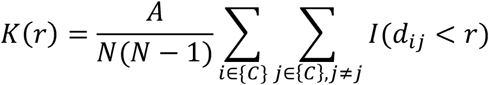

In this equation, *A* represents the nucleus area for each cell, *N* is the total number of CENPC spots in the nucleus, *d*_*ij*_ stands for the Euclidean distance between the i-th and j-th spots, and *r* is a predefined radius within which we evaluate the clustering. The indicator function, *I*(), returns 1 if (*d*_*ij*_ < *r*), and 0 otherwise. The calculation of K(r) involves the summation over all unique pairs of points (*i, j*) in the cell (C). The resulting sum is then normalized by multiplying it to the ratio between A and the product of N and N-1.

To correct for edge effects, which can potentially bias results for spots in proximity of the nucleus ROI periphery, we employ Ripley’s edge-corrected K-function. The correction to the K-function adds a weighting term for each point that is inversely proportional to the area of the region accessible to other points within the specified radius, *r*, without crossing the boundary of the study area. This results in a correction factor that adjusts for the reduced probability of finding neighboring points near the edges of the region under study. The K(r) calculations were performed using the Astropy package in a Jupyter notebook separate from HiTIPS. Finally, the difference between a Poisson point process (representing complete spatial randomness) and the actual data from the cell is computed. The percentage of radii where the measured value of K(r) is higher than the K(r) for the Poisson process is then calculated as a clustering score on a per cell basis.

### Statistical Analysis

Statistical analysis for the DNA FISH and for the CENPC clustering data was performed using the R statistical programming language, and these R packages: tidyverse, data.table, fs, and ggthemes.

Statistical analysis for the MS2-GFP live cell data was performed in Python 3.9 using these libraries: pandas, Seaborn and Matplotlib.

## Acknowledgements

We would like to thank all the members of the Misteli and Larson laboratories for insightful discussions on high-throughput imaging and automated image analysis. This work utilized the computational resources of the NIH HPC Biowulf cluster (http://hpc.nih.gov). We would like to thank the NIH HPC group for their help with data management and software package management. Research in the Misteli Lab, Larson Lab, and HiTIF was supported by the Intramural Research Program of the NIH, NCI, Center for Cancer Research via 1-ZIA-BC010309-24, 1-ZIA-BC011383-12, and 1-ZIC-BC011567-09, respectively.

## Author Contributions

AK and GP established the requirements for HiTIPS. AK wrote all the HiTIPS code base. FA, KG, and VS performed cell culture and treated cells for DNA FISH, IF, and live cell imaging, respectively. FA, KG, and VS acquired the images with high-throughput microscopes. AK, FA, and KG analyzed the high-throughput imaging datasets using HiTIPS and performed statistical analysis and plotting. AK, FA, KG, VS, NF, CHB, DRL, and TM, and GP provided guidance and feedback on the algorithms for image analysis and on the design of the graphical user interface. AK and GP wrote the manuscript. All authors edited and approved the manuscript.

## Code and Data Availability

The HiTIPS code base can be found here: https://github.com/CBIIT/HiTIPS.

## References

1. Boutros, M., Heigwer, F. & Laufer, C. Microscopy-Based High-Content Screening. Cell 163, 1314–1325 (2015).

2. Pegoraro, G. & Misteli, T. High-Throughput Imaging for the Discovery of Cellular Mechanisms of Disease. Trends Genet 33, 604–615 (2017).

3. Joyce, E. F., Williams, B. R., Xie, T. & Wu, C.-T. Identification of genes that promote or antagonize somatic homolog pairing using a high-throughput FISH-based screen. PLoS Genet 8, e1002667 (2012).

4. Shachar, S., Voss, T. C., Pegoraro, G., Sciascia, N. & Misteli, T. Identification of Gene Positioning Factors Using High-Throughput Imaging Mapping. Cell 162, 911–923 (2015).

5. Jowhar, Z. et al. Effects of human sex chromosome dosage on spatial chromosome organization. Mol Biol Cell 29, 2458–2469 (2018).

6. Finn, E. H. et al. Extensive Heterogeneity and Intrinsic Variation in Spatial Genome Organization. Cell 176, 1502–1515.e10 (2019).

7. Park, D. S. et al. High-throughput Oligopaint screen identifies druggable 3D genome regulators. Nature (2023) doi:10.1038/s41586-023-06340-w.

8. Neumann, B. et al. Phenotypic profiling of the human genome by time-lapse microscopy reveals cell division genes. Nature 464, 721–727 (2010).

9. Cuylen, S. et al. Ki-67 acts as a biological surfactant to disperse mitotic chromosomes. Nature 535, 308–312 (2016).

10. Kubben, N. et al. Repression of the Antioxidant NRF2 Pathway in Premature Aging. Cell 165, 1361–1374 (2016).

11. Samwer, M. et al. DNA Cross-Bridging Shapes a Single Nucleus from a Set of Mitotic Chromosomes. Cell 170, 956–972.e23 (2017).

12. Jevtić, P. et al. The nucleoporin ELYS regulates nuclear size by controlling NPC number and nuclear import capacity. EMBO Rep 20, (2019).

13. Schibler, A. C., Jevtic, P., Pegoraro, G., Levy, D. L. & Misteli, T. Identification of epigenetic modulators as determinants of nuclear size and shape. Elife 12, (2023).

14. Stavreva, D. A. et al. Transcriptional Bursting and Co-bursting Regulation by Steroid Hormone Release Pattern and Transcription Factor Mobility. Mol Cell 75, 1161–1177.e11 (2019).

15. Wan, Y. et al. Dynamic imaging of nascent RNA reveals general principles of transcription dynamics and stochastic splice site selection. Cell 184, 2878–2895.e20 (2021).

16. Ljosa, V. & Carpenter, A. E. Introduction to the quantitative analysis of two-dimensional fluorescence microscopy images for cell-based screening. PLoS Comput Biol 5, e1000603 (2009).

17. Imbert, A. et al. FISH-quant v2: a scalable and modular tool for smFISH image analysis. RNA N. Y. N 28, 786–795 (2022).

18. Carpenter, A. E. et al. CellProfiler: image analysis software for identifying and quantifying cell phenotypes. Genome Biol. 7, R100 (2006).

19. Linkert, M. et al. Metadata matters: access to image data in the real world. J. Cell Biol. 189, 777–782 (2010).

20. Moore, J. et al. OME-Zarr: a cloud-optimized bioimaging file format with international community support. bioRxiv 2023.02.17.528834 (2023).

21. Misteli, T. The Self-Organizing Genome: Principles of Genome Architecture and Function. Cell 183, 28–45 (2020).

22. Lieberman-Aiden, E. et al. Comprehensive Mapping of Long-Range Interactions Reveals Folding Principles of the Human Genome. Science 326, 289–293 (2009).

23. Rao, S. S. P. et al. Cohesin Loss Eliminates All Loop Domains. Cell 171, 305_ik_20.e24 (2017).

24. Luppino, J. M. et al. Cohesin promotes stochastic domain intermingling to ensure proper regulation of boundary-proximal genes. Nat Genet 52, 840–848 (2020).

25. Natsume, T., Kiyomitsu, T., Saga, Y. & Kanemaki, M. T. Rapid Protein Depletion in Human Cells by Auxin-Inducible Degron Tagging with Short Homology Donors. Cell Rep 15, 210–218 (2016).

26. Przewloka, M. R. et al. CENP-C is a structural platform for kinetochore assembly. Curr. Biol. CB 21, 399–405 (2011).

27. Foley, E. A. & Kapoor, T. M. Microtubule attachment and spindle assembly checkpoint signalling at the kinetochore. Nat. Rev. Mol. Cell Biol. 14, 25–37 (2013).

28. Muller, H., Gil, J. & Drinnenberg, I. A. The Impact of Centromeres on Spatial Genome Architecture. Trends Genet. TIG 35, 565–578 (2019).

29. Hoencamp, C. et al. 3D genomics across the tree of life reveals condensin II as a determinant of architecture type. Science 372, 984–989 (2021).

30. Kiskowski, M. A., Hancock, J. F. & Kenworthy, A. K. On the Use of Ripley’s K-Function and Its Derivatives to Analyze Domain Size. Biophys. J. 97, 1095–1103 (2009).

31. Rodriguez, J. et al. Intrinsic Dynamics of a Human Gene Reveal the Basis of Expression Heterogeneity. Cell 176, 213–226.e18 (2019).

32. Carpenter, A. E. et al. CellProfiler: image analysis software for identifying and quantifying cell phenotypes. Genome Biol 7, R100 (2006).

33. Stirling, D. R. et al. CellProfiler 4: improvements in speed, utility and usability. BMC Bioinformatics 22, 433 (2021).

34. Hart, T. et al. High-Resolution CRISPR Screens Reveal Fitness Genes and Genotype-Specific Cancer Liabilities. Cell 163, 1515–1526 (2015).

35. Finn, E., Misteli, T. & Pegoraro, G. High-Throughput DNA FISH (hiFISH). Methods Mol Biol 2532, 245–274 (2022).

36. Coulon, A. et al. Kinetic competition during the transcription cycle results in stochastic RNA processing. Elife 3, (2014).

37. Bannon, D. et al. DeepCell 2.0: Automated cloud deployment of deep learning models for large-scale cellular image analysis. 505032 Preprint at 10.1101/505032 (2018).

38. Stringer, C., Wang, T., Michaelos, M. & Pachitariu, M. Cellpose: a generalist algorithm for cellular segmentation. Nat. Methods 18, 100–106 (2021).

39. Hollandi, R. et al. Nucleus segmentation: towards automated solutions. Trends Cell Biol. 32, 295–310 (2022).

40. Stringer, C., Wang, T., Michaelos, M. & Pachitariu, M. Cellpose: a generalist algorithm for cellular segmentation. Nat. Methods 18, 100–106 (2021).

41. Pachitariu, M. & Stringer, C. Cellpose 2.0: how to train your own model. Nat Methods 19, 1634–1641 (2022).

42. Greenwald, N. F. et al. Whole-cell segmentation of tissue images with human-level performance using large-scale data annotation and deep learning. Nat. Biotechnol. 40, 555–565 (2022).

43. Yang, X., Li, H. & Zhou, X. Nuclei Segmentation Using Marker-Controlled Watershed, Tracking Using Mean-Shift, and Kalman Filter in Time-Lapse Microscopy. IEEE Trans. Circuits Syst. Regul. Pap. 53, 2405–2414 (2006).

44. Soille, P. Morphological Image Analysis. (Springer, 2004). doi:10.1007/978ik-662-05088-0.

45. Ulicna, K., Vallardi, G., Charras, G. & Lowe, A. R. Automated Deep Lineage Tree Analysis Using a Bayesian Single Cell Tracking Approach. Front. Comput. Sci. 3, (2021).

46. Kalman, R. E. A New Approach to Linear Filtering and Prediction Problems. J. Basic Eng. 82, 35–45 (1960).

47. Moen, E. et al. Accurate cell tracking and lineage construction in live-cell imaging experiments with deep learning. 803205 Preprint at 10.1101/803205 (2019).

48. Kuhn, H. W. The Hungarian method for the assignment problem. Nav. Res. Logist. Q. 2, 83–97 (1955).

49. Lowe, D. G. Distinctive Image Features from Scale-Invariant Keypoints. Int. J. Comput. Vis. 60, 91–110 (2004).

50. Bay, H., Ess, A., Tuytelaars, T. & Van Gool, L. Speeded-Up Robust Features (SURF). Comput. Vis. Image Underst. 110, 346–359 (2008).

51. Ma, W. et al. Remote Sensing Image Registration With Modified SIFT and Enhanced Feature Matching. IEEE Geosci. Remote Sens. Lett. 14, 3–7 (2017).

52. Thompson, R. E., Larson, D. R. & Webb, W. W. Precise Nanometer Localization Analysis for Individual Fluorescent Probes. Biophys. J. 82, 2775–2783 (2002).

